# High Sucrose Diets Contribute to Brain Angiopathy with Impaired Glucose Uptake, and Psychosis-related Higher Brain Dysfunctions in Mice

**DOI:** 10.1101/2020.02.14.939546

**Authors:** Shinobu Hirai, Hideki Miwa, Tomoko Tanaka, Kazuya Toriumi, Yasuto Kunii, Hiroko Shimbo, Takuya Sakamoto, Mizuki Hino, Ryuta Izumi, Atsuko Nagaoka, Hirooki Yabe, Tomoya Nakamachi, Seiji Shioda, Takashi Dan, Toshio Miyata, Yasumasa Nishito, Kazuhiro Suzuki, Mitsuhiro Miyashita, Toshifumi Tomoda, Takatoshi Hikida, Junjiro Horiuchi, Masanari Itokawa, Makoto Arai, Haruo Okado

## Abstract

Metabolic dysfunction is thought to contribute to the severity of psychiatric disorders; however, it has been unclear whether current high-simple-sugar diets contribute to pathogenesis of these diseases. Here we demonstrate that a high-sucrose diet during adolescence induces psychosis-related behavioral endophenotypes, including hyperactivity, poor working memory, impaired sensory gating, and disrupted interneuron function in mice deficient for glyoxalase-1 (Glo1), an enzyme involved in detoxification of sucrose metabolites. Further, the high-sucrose diet induced microcapillary impairments and reduced brain glucose uptake in brains of *Glo1* deficient mice. Aspirin protected against this angiopathy, enhancing brain glucose uptake, and preventing abnormal behavioral phenotypes. Similar vascular damage to our model mice was found in the brains of randomly collected schizophrenia and bipolar disorder patients, suggesting that psychiatric disorders are associated with angiopathy in the brain caused by various environmental stresses, including metabolic stress.

## Introduction

Considering the global increase in consumption of simple sugars, the World Health Organization published guidelines that addressed concerns regarding body weight gain and dental caries development in 1995 (*1*). High-sugar intake alone increases the risks of numerous chronic diseases, including diabetes, hypertension, and kidney disease. However, there are few studies on the effects of high-sugar intake during adolescence on future mental health. The daily caloric intake from simple sugar by teenagers is higher than that observed for other age groups (∼20% of total daily caloric intake) (*2*). Most chronic psychiatric disorders, including schizophrenia (SZ) and bipolar disorder (BD) discussed in the present study, develop before the age of 30 via complex interactions among multiple genetic and environmental risk factors. Interestingly, patients with SZ and BD consume approximately 2-fold more sugar than age-matched healthy individuals, and patients with SZ who consume more sucrose exhibit more severe symptoms (*3–5*). Moreover, the odds ratios for mental distress, hyperactivity, and behavioral disorders were the highest among adolescents who self-reported the highest consumption of soft drinks (*6*).

Advanced glycation end products (AGEs) and reactive carbonyl compounds are produced from irreversible non-enzymatic crosslinking reactions between carbonyl-containing molecules (e.g. sugars, reactive carbonyl compounds) and other molecules (e.g. amino groups of proteins), and AGEs can lead to the induction of inflammation and oxidative stress in various tissues, leading to a positive feedback loop that further increases AGEs (*7*). There is substantial evidence that patients with psychiatric disorders have elevated proinflammatory cytokines (e.g. interleukin (IL)-1β, IL-6) and increased oxidative stress (*8–11*). Glyoxalase I (GLO1) is a zinc metalloenzyme that protects cells from AGE toxicity by catalyzing the reaction between the reactive carbonyl compound methylglyoxal to glutathione to form S-lactoyl-glutathione (*12*). Patients with depressive-state BD and major depressive disorder have reduced amounts of GLO1, suggesting that this reduction, in combination with an increase in sugar consumption, may be a cause of the increased inflammation and oxidative stress observed in these diseases (*13*). Furthermore, a SZ patient exhibiting poor convalescence is found to harbor a frameshift mutation in *GLO1* leading to reduced enzyme activity (*14, 15*).

Despite accumulating evidence, it is still unproven that excessive sugar intake contributes to the pathogenesis of psychiatric disorders among susceptible individuals. We addressed this question by generating mice with psychosis-related phenotypes associated the gene × environment interactions (G × E) to determine whether excessive sucrose consumption during adolescence is a novel environmental risk factor for SZ and BD. We further identified angiopathy in mice with high-sucrose intake and genetic venerability, and confirmed similar vascular damage in randomly collected brain samples from BD and SZ patients. This suggests that a variety of genetic and environmental factors can adversely affect brain capillaries. In addition, we found that chronic anti-inflammatory drug treatment prevented core phenotypes of psychiatric disorders as well as vascular damage, suggesting that angiopathy may have an important influence on the development and/or condition of psychiatric disorders.

## Results

### Psychosis-related behavioral phenotypes in mice on a high-sucrose diet

We fed mice one of two diets containing the same total calories and caloric proportions of carbohydrates, fat, and proteins, but with either starch or sucrose as the main carbohydrate (Fig. 1A). We investigated four groups of mice fed these diets for 50 days immediately after weaning (from postnatal day 21, corresponding to the juvenile/adolescent stage): starch-fed wild-type (WT) mice (control, CTL), sucrose-fed WT mice (environmental stressor, Env), starch-fed *Glo1* heterozygous knockout (*Glo1/+*) mice (genetic factor, Gen), and sucrose-fed *Glo1/+* mice (G × E) (Fig. 1A). Reduced Glo1 expression in *Glo1/+* mice was confirmed in the cerebral cortices, including the hippocampus, using western blotting (fig. S1A, B). The body weight trajectories of these mice were similar to those of control mice for at least 11 weeks of age, indicative of normal structural development (fig. S1C) and removing obesity as a possible cause of the observed group differences described below. We tested eight different behaviors in mice, including open-field locomotor activity (Fig. 1B), acoustic startle response (Fig. 1C), pre-pulse inhibition (PPI; a measure of sensory-motor gating) (Fig. 1D), object location performance (used as a test of working memory) (Fig. 1E), self-grooming (Fig. 1F), nest building (Fig. 1G), elevated plus maze activity (fig. S2A), and social interaction (fig. S2B). These behaviors could be classified into those that are not affected by either diet or genotype, those that are affected by genotype but not diet, those affected by diet but not genotype, and those affected by the combination of diet and genotype. Thus, neither diet nor genotype, nor a combination of diet and genotype had any significant effect on social interactions (fig. S2B), while acoustic startle responses were reduced in *Glo1/+* mice but not significantly affected by feeding type (Fig. 1C). Self-grooming (Fig. 1F), nest building (Fig. 1G), and elevated plus maze activity (fig. S2A) were affected by diet, but not genotype, while open field locomotor activity, PPI, and object location performance required a combination of sucrose feeding and a *Glo1/+* genotype to be affected. In the open field test (Fig. 1B), neither starch-fed *Glo1/+* mice, nor sucrose-fed WT mice, showed significant differences in distance traveled compared to starch-fed WT mice, while sucrose-fed *Glo1/+* mice showed significant increases. Similarly, starch-fed *Glo1/+* mice and sucrose-fed WT mice had similar PPI scores compared to starch-fed WT mice, while sucrose-fed *Glo1/+* mice showed significant decreases (Fig. 1D). Sucrose-fed WT mice had a non-significant reduction in object location test scores, and this reduction became significant in sucrose-fed *Glo1/+* mice (Fig. 1E). Altogether, our results suggest that a combination of sucrose feeding and *Glo1/+* mutation can generate novel phenotypes reminiscent of psychiatric disease phenotypes that are not found by altering diet or genotype alone. To further support our idea that synthetic G × E interactions may induce or enhance the severity of psychosis-like symptoms, we examined the effects of the high sucrose diet on an additional major psychosis-associated mutation in *Disrupted-in-schizophrenia 1* (*DISC1*) locus (*16*) (fig. S3). DISC1 plays important roles in neurodevelopment and synaptic activity, but a loss-of-function mutation in *DISC1* gene alone is not sufficient to cause psychiatric disorders (*17*). Similar to the results for *Glo1/+* mice, *Disc1/+* mice fed a high sucrose diet (G × E mice) showed severely impaired object location performance (working memory) and PPI (fig. S3C, D), whereas WT mice fed a high sucrose diet or *Disc1/+* mice fed control diet did not, again suggesting that a combination of genetic and environmental influences can induce psychosis-like behaviors.

**Fig. 1.**
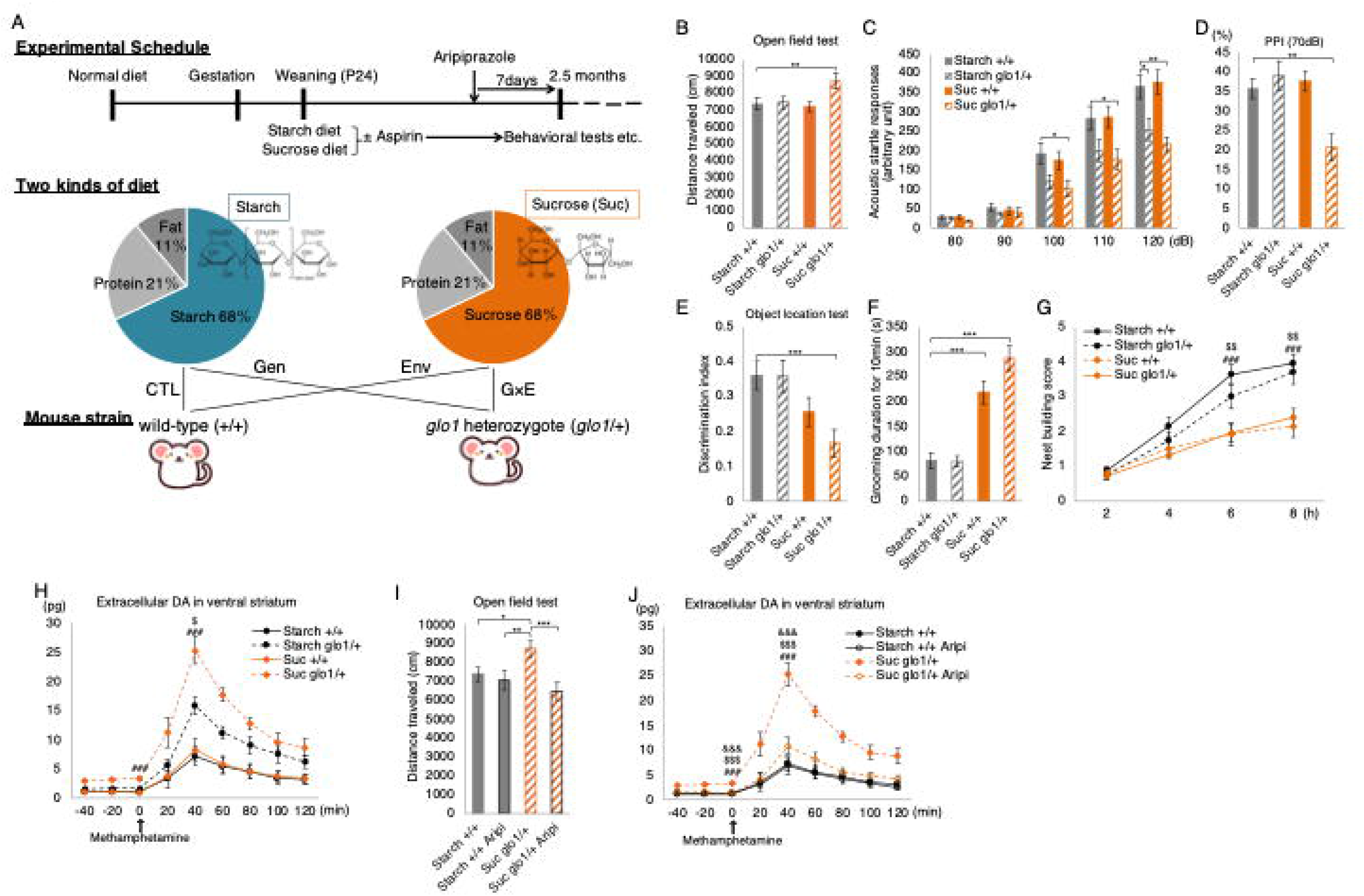
Generation of G × E model mice and psychosis-related phenotype analyses. (**A**) Experimental Timeline. See Materials and Methods for a complete description. **(B–G)** Behavioral analyses in the four groups of mice (n = 18–23 mice per group). (**B**) Spontaneous locomotor activity in the open-field test. (**C**) Acoustic startle responses. (**D**) Pre-pulse inhibition (PPI) using a 70 dB pre-pulse. (**E**) Object location test to evaluate working memory. (**F**) Duration of self-grooming in the home cage. (**G**) Quantification of nest-building skills over 8 h. Post hoc Dunnett‘s multiple comparisons test among groups at each time point indicated ###*p* < 0.001 for Suc *glo1*/+ vs. Ctrl (Starch +/+), $$*p* < 0.01 for Suc +/+ vs. Ctrl (Starch +/+). (**H, J**) Extracellular dopamine (DA) concentration in the nucleus accumbens measured at 20 min intervals using an *in vivo* microdialysis system in the presence or absence of aripiprazole (Aripi). Methamphetamine (1.25 mg/kg) was administered via intraperitoneal (i.p.) injection at time 0 (arrow) (n = 6–11 mice per group). See Materials and Methods for detailed statistical analyses. (**I**) Effects of Aripi treatment on locomotor activity (n = 12–18 mice per group). (**B–H**) Starch +/+ group was used as a control for post hoc Dunnett’s test. All data are presented as mean ± SEM. \*\*\**p* < 0.001, \*\**p* < 0.01, \**p* < 0.05

Schizophrenia (SZ) is associated with increased amphetamine-induced dopamine release in the striatum (*18, 19*) and unbalanced dopamine regulation between the medial prefrontal cortex (mPFC) and striatum (*20*). To examine the effects of high sucrose diet on dopamine release in *Glo1/+* mice, we next measured dopamine levels in the nucleus accumbens (NAc) and mPFC of these mice using *in vivo* microdialysis. We found that the high sucrose diet increased both basal and amphetamine-induced dopamine release in the NAc specifically in *Glo1/+* mice (Fig. 1H), while reducing basal dopamine release in the mPFC in either genetic background (fig. S2C). To assess whether enhanced dopamine release in the NAc could induce the observed behavioral phenotypes, we examined the effects of aripiprazole, a D2 receptor partial agonist and clinical antipsychotic (*21*), administered during the last 7 days (0.5 mg/kg/day) of sucrose feeding (Fig. 1A, I, J and fig. S2D–I). We found that the hyper-locomotion and increased striatal dopamine release in G × E mice were completely reversed by aripiprazole treatment (Fig. 1I, J), while other behavioral defects were not improved (fig. S2D–I). Thus, aripiprazole treatment selectively improves a subset of behavioral abnormalities in G × E mice that are likely caused by dysregulated dopamine signaling.

### Dysfunction of parvalbumin-positive inhibitory interneurons in G × E mice

A reduction in density of parvalbumin (PV)-positive GABAergic interneurons is reported for schizophrenia (*22–25*), and the precisely coordinated activity of these neurons is crucial for the regulation of PPI and working memory (*26*). Thus, we next investigated PV expression using immunohistochemistry and western blotting to determine whether alterations in PV signaling may contribute to our observed psychiatric disease-associated phenotypes. A high sucrose diet caused a reduction in PV-positive cells in both *Glo1/+* and WT mice (Fig. 2A, B) and significantly reduced PV expression in sucrose-fed *Glo1/+* mice (Fig. 2C, D), suggesting that the sucrose diet and the *Glo1*/+ mutation had additive or synergistic effects on PV signaling. Since gamma oscillations are produced via the synchronous activation of PV neurons, we measured gamma oscillations (30–45 Hz) using surface electroencephalography (EEG) to determine whether downregulation of PV protein was accompanied by functional abnormalities in neural activity. Interestingly, sucrose-fed mice (WT and *Glo1*/+ mice) exhibited elevated baseline gamma oscillation power compared with control mice (starch-fed WT) in the home cage (Fig. 2E). Further, only sucrose-fed *Glo1*/+ mice failed to exhibit an increase in the gamma oscillation power when approaching a novel object (Fig. 2F). These results are consistent with findings in patients with SZ and BD as well as other mouse models of psychosis demonstrating increased baseline gamma oscillations and decreased sensory stimulus-evoked synchronized gamma power (*27, 28*). Thus, our results suggest that G × E mice mimic the pathophysiological changes of PV neurons observed in psychiatric disorders.

**Fig. 2.**
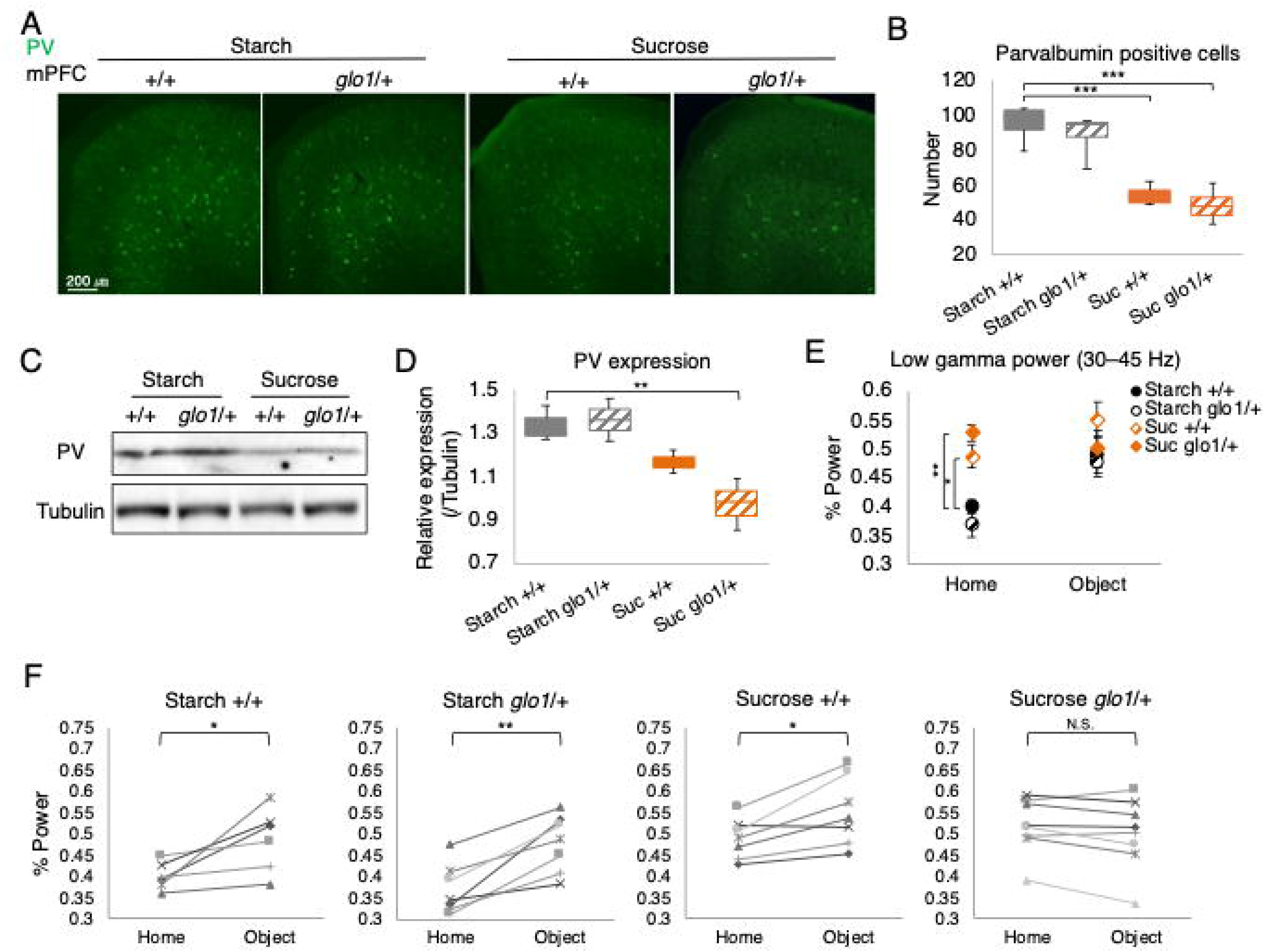
Parvalbumin-positive interneuron dysfunction in G × E mice. (**A**) Parvalbumin (PV) immunohistochemistry in the medial prefrontal cortex (mPFC). (**B**) Number of PV-positive cells in the mPFC (n = 4 mice per group). (**C**) Western blot analysis of PV protein expression using tubulin as internal control. (**D**) Densitometric analysis of PV protein expression: PV band intensities were divided by the corresponding tubulin band intensities (n = 3 mice per group). (**E**) Average gamma band power in the home cage and during novel object recognition in the open field (n = 7–8 mice per group). (**F**) Changes in gamma band power from the home cage to the novel object phase in individual mice (n = 7–8 mice per group). Each *p* value indicates the result of repeated measures analysis of variance (ANOVA). The effect of changes in gamma power at Starch +/+ (F1, 5 = 8.29, *p* = 0.035), at Starch *Glo1*/+ (F1, 6 = 29.75, *p* = 0.0016), at Sucrose *+*/+ (F1, 6 = 12.04, *p* = 0.013), at Sucrose *Glo1*/+ (F1, 7 = 4.038, *p* = 0.084). (B, D, H) Starch +/+ group was used as control for post hoc Dunnett’s test. All data are presented as mean ± SEM. \*\*\**p* < 0.001, \*\**p* < 0.01, \**p* < 0.05

To summarize, the administration of a high-sucrose diet to *Glo1/+* mice induces behavioral, histological, and pathophysiological phenotypes reminiscent of phenotypes observed in psychiatric disorders. This suggests that excessive sucrose intake during adolescence is a potential environmental risk factor for these diseases.

### AGE accumulation and impaired astrocyte function in G × E mice

How does a decrease in GLO1 expression and an increase in sucrose consumption lead to defects in activity of PV and dopaminergic neurons? We first examined GLO1 expression and detected high expression in astrocytes (fig. S4A–G), especially those surrounding capillaries (fig. S4B, C), moderate expression in neurons (fig. S4H, I), and low expression (below detection limits) in vascular endothelial cells (fig. S4A–G) and microglial cells (fig. S4H, J). Expression was not detected in the brains of *Glo1* homozygous mice (fig. S3K) indicating that the antibody we used was specific for GLO1.

Since both GLO1 and a high sucrose diet are associated with glycation toxicity, we next examined AGE immunoreactivity to detect levels of AGE products of glucose metabolism and/or chemical reaction of glucose. We found stronger AGE immunoreactive signals in the vascular endothelial cells of G × E mice compared with control mice (Fig. 3A–H). Moreover, we detected AGE products of methylglyoxal metabolism and/or chemical reaction of methylglyoxal using AGE-4 antibody, in the microglia of sucrose-fed mice (fig. S5A–E). Compared to controls, AGE accumulation in sucrose-fed *Glo1/+* mice was accompanied by elevated IBA1 immunofluorescence intensity and an enlarged CD68-positive area (fig. S5F, G, H), both of which are phenotypes identified in activated microglia (*29, 30*). Thus, sucrose feeding in *Glo1/+* mice results in microglial and endothelial AGE accumulation and microglial activation.

**Fig. 3.**
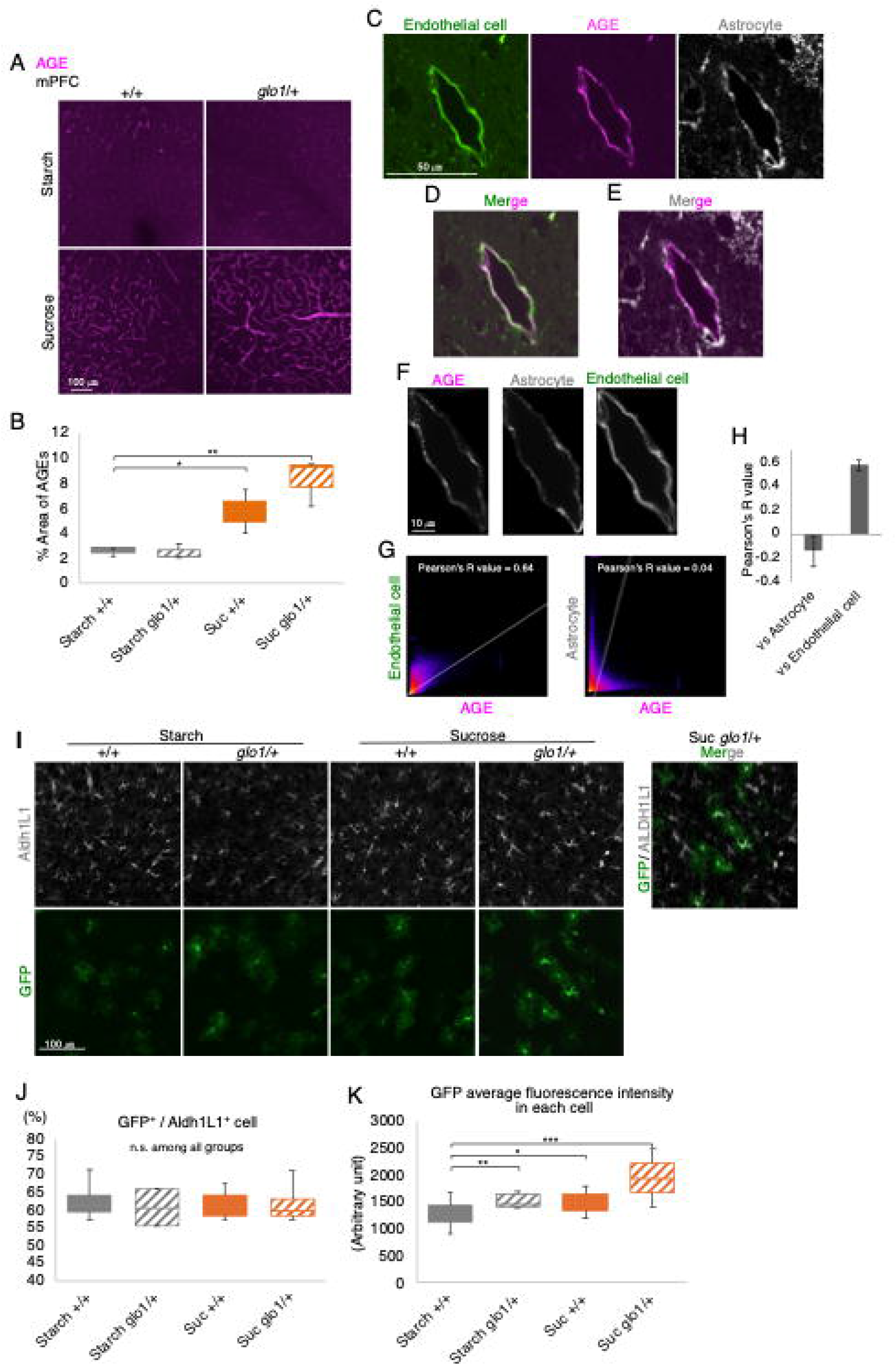
AGE accumulation in the neurovascular endothelium and pre-inflammatory status of astrocytes in G × E mice. (**A**) AGE immunohistochemistry in the medial prefrontal cortex. (**B**) Measurement of AGE immunoreactive area. The mean intensity of the AGE-immunopositive area in the entire image was measured for each section (n = 3 mice per group). (**C–E**) Colocalization of the endothelial cell marker tomato lectin with the astrocytic marker ALDH1L1 or AGEs. (**F**) Enlarged version of images in (**C**) presenting the region used for colocalization analysis. 2D intensity histogram of the two indicated channels for identifying the colocalization of AGE with ALDH1L1 (Astrocyte) or tomato lectin (Endothelial cell). (**H**) Average R-value of colocalization data including (**G**) three different locations of three mice. (**I**) Immunohistological images of GFP-positive astrocytes and ALDH1L1 in the mPFC region and merged GFP/ALDH1L1 image. (**J**) Percentage of GFP-positive cells per total ALDH1L1-positive cells in each image in (**I**). (**K**) Mean fluorescent GFP intensities of 10 randomly selected cells per image in (**I**) (from four independent mice). (**B, J, K**) The Starch +/+ group was used as control for post hoc Dunnett’s test. All data are presented as mean ± SEM. \*\*\**p* < 0.001, \*\**p* < 0.01, \**p* < 0.05

Since astrocytes express Glo1 most strongly (fig. S4A–G), we next used mice expressing green fluorescent protein (GFP) under control of the glial fibrillary acid protein (GFAP) promoter (*31, 32*) to evaluate how astrocytes are affected in our G x E condition. Typically, cellular damage from AGEs is caused by inflammatory responses induced by Receptor of AGEs (RAGE) activation or by a loss of normal protein function following AGE-forming reactions (*33*). Astrocytes exhibit a well-described reactive phenotype in response to pathogenic conditions including neuroinflammation characterized by enhanced GFAP expression (*34, 35*). Strongly enhanced *GFAP* promoter function was observed in G × E mice, without changes in the number of GFAP-positive astrocytes (Fig. 3I–K), indicating that the astrocytes in G × E mice are in the reactive pre-condition during the high-sucrose feeding. Taken together, these results suggest that a high sucrose diet enhances AGE production in endothelial cells and microglia (Fig. 3A–H and fig. S5A–E), and converts astrocytes into a pre-inflammatory state in *Glo1*-deficient mice (Fig. 3I–K).

### Microcapillary angiopathy and impaired glucose intake in G × E mice

Endothelial cells and astrocytes are functional and physical components of the blood-brain barrier (BBB) which tightly controls the parenchymal environment by modulating the selective passage of nutrients and various factors (*36*). Thus, endothelial AGE accumulation and astrocyte reactivity may impair BBB function. To examine changes in endothelial function, we first conducted a transcriptome analysis of the prefrontal cortex (PFC), a region strongly implicated in psychiatric impairments, using microarrays (Fig. 4A, B). The coagulation factor V, essential for the production of fibrin from fibrinogen, ranked seventh on a list of transcripts exhibiting more than doubled expression in G × E mice compared with the other three groups (Fig. 4A, Supplementary Table 1, 2). Fibrin controls hemostasis via polymerization with platelets to form blood clots, and deposits of this protein are indicative of endothelial abnormalities (*37*). In early stages of endothelial cell impairment, fibrin accumulates in capillaries. Therefore, we investigated vascular fibrin accumulation using immunohistochemistry and confirmed the presence of significant fibrin accumulation on the vascular lumen side of endothelial cells in brain capillaries of G × E mice (Fig. 4C–G). Fibrin leakage and deposition in the brain parenchyma, as observed in Alzheimer’s disease, was not detected in G × E mice, indicating that physical BBB disruption did not occur (*38*).

**Fig. 4.**
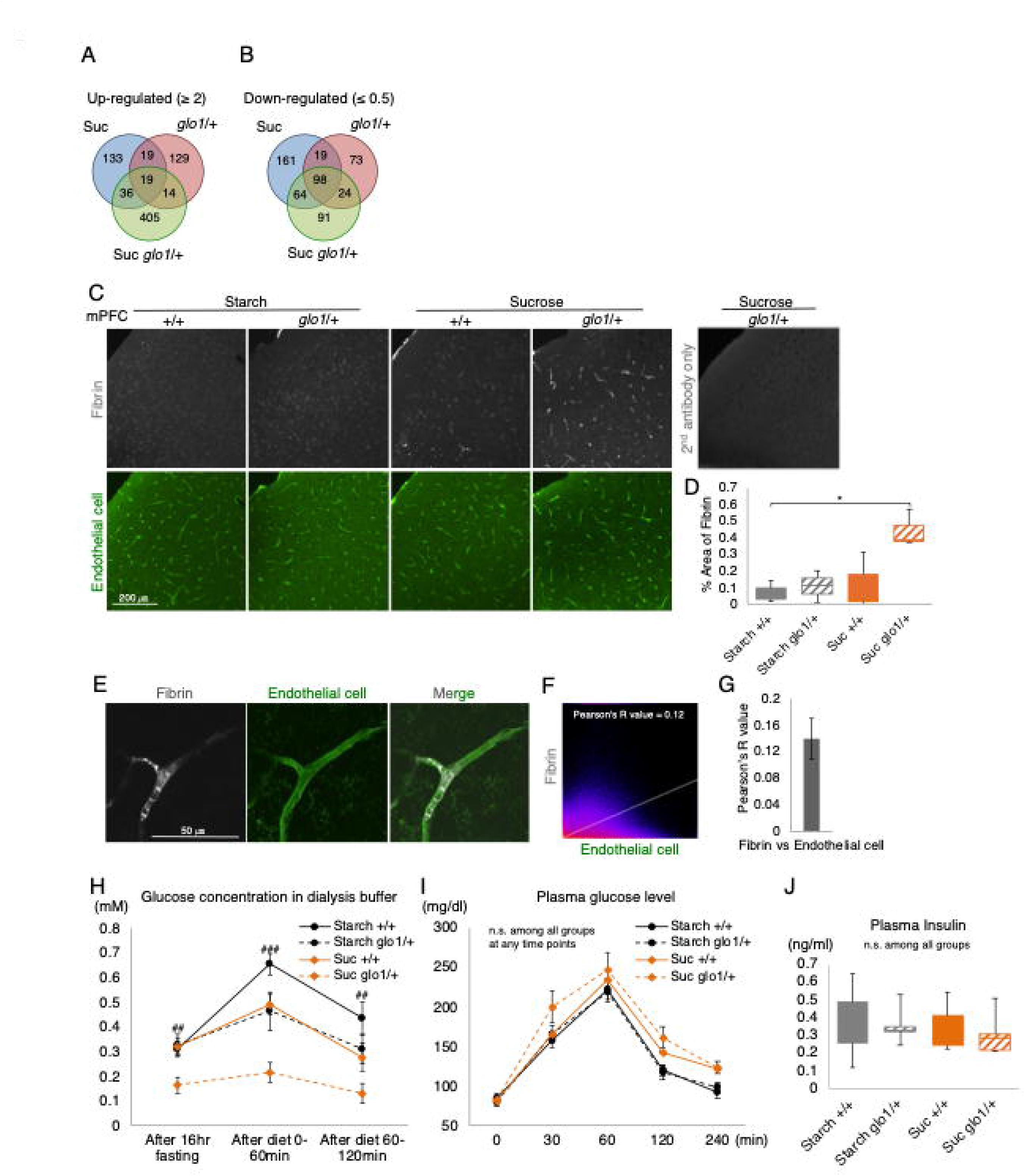
Angiopathy and impaired glucose transport in G × E mice. (**A, B**) Venn graph showing overlap in prefrontal cortex genes exhibiting a >2-fold (**A**) or <0.5-fold (**B**) expression change compared with the control (CTL) group (n = 3 mice per group). (**C**) Immunohistochemical images of fibrin and the endothelial cell marker tomato lectin. (**D**) Measurement of the area of fibrin immunoexpression in (**C**). The mean intensity of the fibrin-immunopositive area of the entire image was measured for each section (n = 4 mice per group). (**E**) Magnified immunohistochemical images of fibrin and the endothelial cell marker tomato lectin. (**F**) 2D intensity histogram of the two indicated channels using the left and middle panels in (**E**) for identifying fibrin colocalization with tomato lectin. (**G**) Average R-value of colocalization data in **(F)** from three different locations of three mice. Extracellular glucose concentrations in the dialysis buffer from mPFC samples at each time point (n = 5–6 mice per group). See the Materials and Methods for detailed statistical analyses. (**I**) Plasma glucose concentrations (n = 6–7 mice per group). The first measurement was performed after 16 h of fasting, and the second blood collection 30 min after eating 0.05 g of carbohydrates. (**J**) Fasting plasma insulin levels (n = 5–6 mice per group). (**D, H**–**J**) Starch +/+ group was used as a control for post hoc Dunnett’s test. All data are presented as mean ± SEM. \**p* < 0.05

We next speculated that the abnormal vascular endothelial cells and astrocytes observed in G × E mice could alter glucose uptake from the plasma into the brain parenchyma. Extracellular glucose concentrations in the brain parenchyma were measured under three conditions: 1) fasting, 2) 1 h after feeding, and 3) 2 h after feeding. We first measured a baseline glucose concentration in the mPFC in starch-fed WT mice after starvation for 16 hrs (Fig. 4H). This concentration increased significantly within an hour after starch feeding, and decreased close to baseline 2 hrs after feeding. Sucrose-fed WT mice and starch-fed *Glo1/+* mice had similar baseline glucose concentrations, similar increases 1 hr after starch or sucrose feeding, and similar decreases back to baseline 2 hrs after feeding. In contrast, sucrose-fed *Glo1/+* mice had both significantly decreased basal glucose concentrations in the mPFC, and no apparent increase in glucose concentrations after sucrose feeding. We measured vascular diameter and expression of glucose transporter 1, the major glucose transporter expressed in vascular endothelial cells and astrocytes, and observed no differences among the four groups (fig. S6). Further, no differences were detected in plasma glucose after starvation and feeding, or in fasting plasma insulin levels among the four groups (Fig. 4I, J), indicating that the lower parenchymal glucose concentrations in G × E mice is due to reduced uptake across the BBB rather than dysregulation of plasma glucose or insulin signaling.

### Protective effects of chronic low-dose aspirin against behavioral abnormalities and angiopathy

Previous reports have shown that adjunct non-steroidal anti-inflammatory drug (NSAID) treatment can improve psychiatric disorder scores (*39, 40*). Aspirin, an NSAID, is routinely used for the prevention and alleviation of vascular-related adverse events associated with high blood pressure, ischemia, and cardiovascular diseases (*41, 42*). Here, we examined whether aspirin treatment can protect against the development of psychiatric phenotypes in G × E mice (Fig. 1A, Fig. 5A–D and fig. S7A–I). Low-dose aspirin (1 mg/kg/day) fully prevented hyperlocomotor activity, deficits in PPI, working memory, grooming duration (Fig. 5A–D), and partially prevented abnormal enhancement of DA release after methamphetamine administration and reduced PV expression among G × E mice (fig. S7D–F), but did not improve acoustic startle responses, nest building and elevated plus maze scores and abnormal astrocyte activation (fig. S7A–C, G–I). These improvements were accompanied by a decrease in endothelial fibrin accumulation (Fig. 5E, F) and a partial restoration of glucose intake into the brain parenchyma (Fig. 5G). Collectively, these results suggest that aspirin treatment in G × E mice significantly improves angiopathy and brain glucose availability, contributing to prevention of several behavioral abnormalities.

**Fig. 5.**
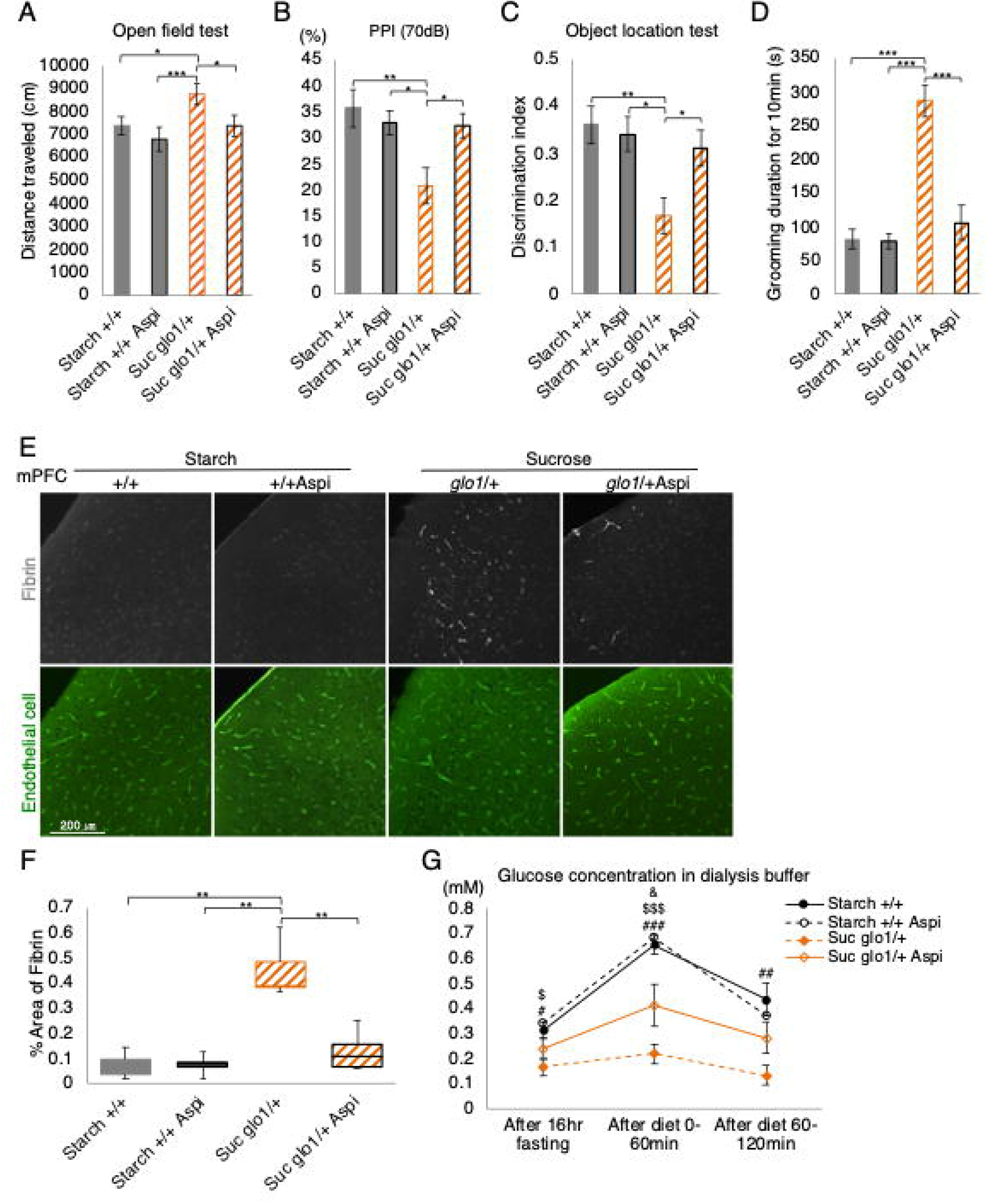
Protective effects of low-dose aspirin in G × E mice. (**A–D**) Results of behavioral tests performed to evaluate the effects of aspirin treatment (n = 12–21 mice per group). (**A**) Quantification of spontaneous locomotor activity in the open field. (**B**) Pre-pulse inhibition at 70 dB. (**C**) Object location test of working memory. (**D**) Quantification of nest-building skills over 8 h. (**E**) Immunohistochemical images of fibrin and the endothelial cell marker tomato lectin. (**F**) Measurement of the area of fibrin immunoexpression in (**E**). The mean intensity of the fibrin-immunopositive area of the entire image was measured for each section (n = 3 mice per group). (**G**) Extracellular glucose concentrations in the dialysis buffer at each time point (n = 4–6 mice per group). Post hoc Tukey‘s multiple comparisons test of groups at each time point, ^###^*p* < 0.001, ^##^*p* < 0.01, ^#^*p* < 0.05 for Starch +/+ vs. Suc *Glo1*/+, ^$$$^*p* < 0.001, ^$^*p* < 0.05 for Starch +/+ Aspi vs. Suc *Glo1*/+, ^&^*p* < 0.05 for Starch +/+ vs. Suc *Glo1*/+ Asp. All data are presented as mean ± SEM. \*\*\**p* < 0.001, \*\**p* < 0.01, \**p* < 0.05

### Angiopathy in postmortem brains of patients with psychiatric disorders

Finally, we compared immunostains of brain slices from healthy controls and patients with SZ or BD to examine whether these patients exhibit angiopathic fibrin accumulation in vascular endothelial cells. Similar to our findings in G × E mice, we found significantly elevated fibrin accumulation in the vascular endothelium of brain slices from randomly collected SZ and BD patients (Fig. 6A–C). Therefore, fibrin-related angiopathy and vascular damage may be a novel and common phenotype of psychiatric illness contributing to disease progression.

**Fig. 6.**
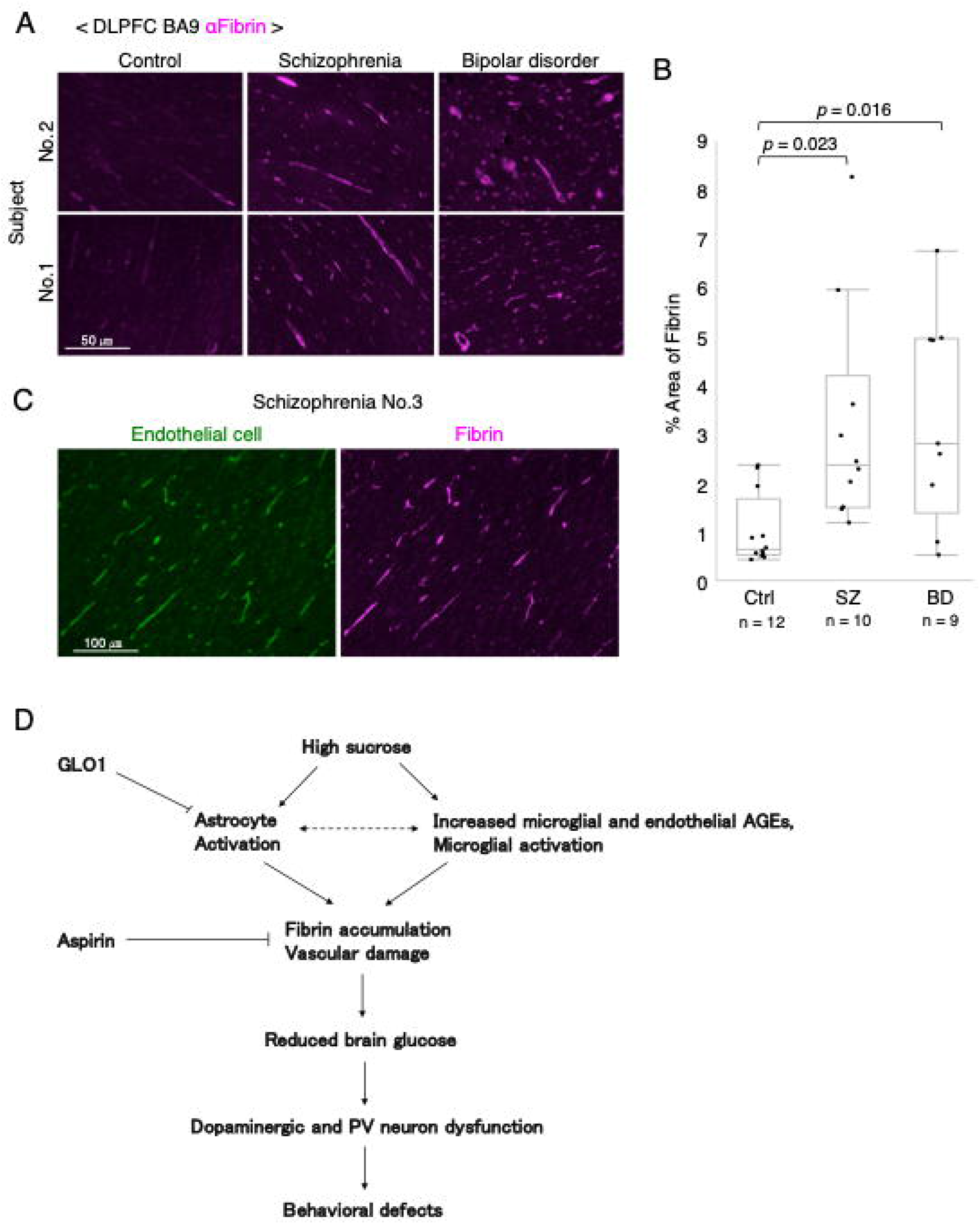
Angiopathy in postmortem brains from individuals with psychiatric disorders. (**A**) Representative immunohistochemical fibrin images in the BA9 region of postmortem brains from controls and patients with schizophrenia (SZ) or bipolar disorder. (**B**) Measurement of the area of fibrin immunoexpression in (**A**). The mean intensity of the fibrin-immunopositive area of the entire image was measured for each section. (**C**) Representative immunohistochemical images of fibrin (magenta) and the endothelial cell marker tomato lectin (green) in postmortem brains from a patient with SZ. Fibrin-positive areas are merged with areas of vascular endothelial cell marker expression. (**D**) Diagrams describing the hypothesis proposed to explain functional and behavioral abnormalities in control (CTL) mice (left) and G × E mice (right) (see the Discussion for details). All data are presented as mean ± SEM. \*\**p* < 0.01

## Discussion

Using a mouse model, we demonstrated that high dietary sucrose consumption during adolescence is a potential risk factor for the development of behavioral phenotypes associated with psychiatric illnesses, such as SZ and BD. These behavioral phenotypes include impaired sensory gating, dysfunctions in PV interneurons and working memory, hyperactivity, and increased basal and stimulus-evoked striatal dopamine release. Second, we identified endothelial fibrin accumulation (“angiopathy”) in both our mouse model and in randomly collected postmortem brains of patients with SZ or BD. We also observed that glucose intake from the plasma into the brain parenchyma was impaired in G × E mouse model. Chronic low-dose aspirin treatment prevented fibrin deposition in the capillaries, improved glucose transport, and reversed several behavioral phenotypes in G × E mice, suggesting angiopathy as a seminal pathogenic event in mental illness.

In Fig. 6D we propose a possible model summarizing our results. A high sucrose diet causes AGE accumulation in microglia and endothelial cells which express low amounts of GLO1 (Fig. 3A–H and fig. S5A–E). Astrocytes are less sensitive to this diet in WT animals because they express high amounts of GLO1 to relieve glycation stress (fig. S4A–G). However, the reduced GLO1 in *Glo1*/+ mice causes astrocytes to become activated by a high sucrose diet (Fig. 3I–K). This activation may be caused directly by glycation stress in astrocytes or may occur as a consequence of increased microglial and endothelial AGEs. Astrocytic activation further increases microglial and endothelial AGEs in a positive feedback loop (*7*). Increased microglial and endothelial AGEs and astrocytic activation may induce fibrin accumulation and damage to the brain vasculature (*33*) (Fig. 4C–G). This results in decreased glucose import into the brain and dopaminergic and PV neuron dysfunction which in turn cause psychosis-associated behavioral defects (Fig. 1B–H, Fig. 2 and Fig. 4H).

Strikingly, administration of the NSAID aspirin, which inhibits inflammation and oxidative stress, protected against the emergence of angiopathy as evidenced by reduced fibrin accumulation, partially restored parenchymal glucose concentrations, and prevention of several psychiatric-related phenotypes (Fig. 5 and fig. S7D–F). The increase in coagulation factor V and the accumulation of fibrin in endothelial cells in G x E mice indicate the presence of some vascular damage (*43*) (Fig.4A–G). Aspirin may prevent this damage through its antiplatelet coagulation and anti-inflammatory properties. Since oxidative stress and chronic inflammation are also common features of psychiatric disorders, aspirin may provide a protective effect against the etiology of these diseases as well (*8, 11, 44, 45*). We have tried to quantify ROSs in our experimental mice, but thus far have not been able to see differences among the four groups (fig. S8), so a more sensitive detection method such as FRET probe for ROS detection may be required.

Sucrose consists of glucose and fructose, when these sugars are ingested at the same time, we find that they generate AGEs in different cell types. Glucose-metabolism-derived AGEs, which can be detected using α-AGE polyclonal antibody, accumulated primarily in vascular endothelial cells (Fig. 3A H), while methylglyoxsal-derived AGEs, recognized by the α-AGE-4 monoclonal antibody accumulated specifically in microglia (fig. S5A–E). Our results suggest that these two cell types may import and metabolize these two sugars differently, but more detailed validation, such as one-cell metabolomic analysis, is required to validate this idea.

Robust PV neuron function is required for PPI, working memory, amphetamine-induced hyper-locomotion, dopamine (or 3,4-dihydroxyphenylacetic acid) release regulation, and gamma oscillation generation, all of which are considered core symptoms of psychiatric disorders (*26*). The PV neuron dysfunction in G × E mice may be caused by the unique properties of these cells. PV-expressing interneurons are the fast-spiking neuron subtype exhibiting a lower input resistance and higher-amplitude rapid after-hyperpolarization than several projection neurons, and the combination of these properties generates higher frequency action potentials compared with those by other neuron types. This frequency of high action potentials in PV neurons can be exploited by their fast spike phenotype to produce fast oscillations (30-100 Hz) in the gamma band of neural oscillations (*46*). To maintain this rapid spiking property, PV neurons require high energy expenditure as evidenced by mitochondrial and cytochrome c oxidase enrichment (*47*). Therefore, reduced glucose within the brain parenchyma may preferentially disturb PV neuron function, resulting in reduced PV expression. Indeed, reduced PV neuron number has been repeatedly reported in postmortem brains of patients with psychiatric disorders such as SZ and BP, and this reduction is proposed to be caused by a decrease in PV mRNA or protein expression in these cells, rather than by a loss of PV neurons themselves (*22–25*). Since PV is a Ca^2+^ binding protein, proper regulation of the expression of this protein is considered to be important for maintaining the plasticity of PV neurons. For example, in PV neurons of the hippocampus and mPFC, the expression of PV protein changes after contextual fear conditioning, exposure to rich environments, and pharmacogenetic activation of PV neurons, suggesting PV expression and plasticity are correlated (*25, 48*). Further, it is known that the inhibitory activity of PV neurons is critical under environmental stress to prevent a sequela of excessive excitatory activity such as oxidative stress and inflammation (*47*). Pathogenic processes might more readily occur during the critical adolescent prodromal period, wherein PV neurons may have increased vulnerability to environmental stresses because of delayed maturation, which is another feature of PV neurons (*49*).

In the present study, we identified capillary angiopathy in both G × E mice and postmortem brains of patients with SZ and BD (Fig. 4C–G and Fig. 6A–C). In our G × E mice, angiopathy was caused by the high AGE production capacity of sucrose combined with a GLO1 deficiency. Since high sugar consumption has been associated with SZ and BD, and decreased GLO1 expression and function have been seen in SZ and BD, similar interactions may be associated with some cases of psychiatric diseases. However, various other environmental stresses and genetic conditions may also converge to induce angiopathy. Several studies have reported that stressors, such as social defeat, isolation, and viral infection, induce vascular defects (*50–52*). These stressors are also risk factors for SZ and BD and induce PV neuron hypofunction (*53*), suggesting that angiopathy may be a common trigger for psychiatric phenotypes.

## Acknowledgments

We thank Sayaka Ogikubo, Yoshie Matsumoto, Haimei Zhang, Minami Murata, Izumi Nohara, Yukiko Shimada, Emiko Hama, Nanako Obata, Mai Hatakenaka, and Chikako Ishida for their contribution to experiments related to this research. We also thank Dr. Tohru Kodama for explaining the procedures for microdialysis and encephalogram recording. We are grateful to Dr. Kenji Tanaka for reviewing the study. We acknowledge Chiaki Watanabe and Hiromi Onuma for coordinating the donations of postmortem brains. Further, we thank Prof. Hideki Chiba for the preparation of postmortem brain samples. We express our gratitude to the families of the deceased individuals for the donations of brain tissue and their time and effort devoted to the consent process and interviews.

## Funding

This work was supported by the Japan Society for the Promotion of Science (KAKENHI grants 18K14832 to S.H., 17K18395 and 19K08033 to H.M., 17K16408 to T.T., and 18H02537 and 18K19383 to H.O.); the Ichiro Kanehara Foundation, Japan Prize Foundation, and Takeda Science Foundation (to S.H.); the Naito Foundation (to H.M.); the Strategic Research Program for Brain Sciences from AMED (grant JP19dm0107107 to H.Y.); and a Grant-in-Aid for Scientific Research on Innovative Areas from the MEXT (JP16H06277 to H.Y.). This research was also supported by KAKENHI Grant Numbers 16H05380 (to M.A.), 18K06977 (to K.T.), 19H03589 (to M.I.), and 18K15354 (to K.S.) as well as AMED Grant Number JP19dm0107088 (to M.I.). This study was also supported by The Kanae Foundation for the Promotion of Medical Science (to K.T.) and The Uehara Memorial Foundation (to M.A.), and the Collaborative Research Program of Institute for Protein Research, Osaka University, ICR-21-3.

## Author contributions

S.H. and H.O. designed the study. S.H. performed and analyzed all experiments. H.M. assisted with the experimental design of EEG recording and analysis of the results and edited the manuscript. T.T. coordinated the EEG recording. Y.K., M.H., R.I., A.N., and H.Y. helped with the design of experiments using human specimens and provided fixed human brain sections. T.S. performed data and image analysis. T.N. and S.S. provided GFAP-GFP mice. T.D. and T.M. generated and provided *Glo1* knockout (KO) mice. K.T. and K.S. backcrossed *Glo1* KO mice to B6J mice for all experiments. Y.N. performed cDNA microarray analysis of gene expression. H.S. assisted with behavioral experiments. M.I., M.A., K.T., K.S., and M.M. provided important suggestions to this study. T. T. and T. H provided critical input on *Disc1-LI* mouse experiments. J.H. edited the manuscript. S.H. generated all figures, tables, and wrote the manuscript. H.O. edited the manuscript and supervised this study.

## Competing interests

The authors declare no competing interests.

## Data and materials availability

All materials used in this paper are available upon request. The accession number for transcriptome analysis data is provided in the Materials and Methods.

## Materials and Methods

### Experimental design (related to Fig.1A)

After weaning (P21), wild-type (WT) and *Glo1* heterozygous mutant mice were fed either a starch diet (control) or a sucrose diet (experimental). Diets were equal in total calories and proportions of calories from carbohydrates, lipids, and proteins. The behavioral test battery was administered starting at 2.5 months of age (upper panel). Middle panel: Macronutrient composition of the two diets. We used *Glo1* heterozygous mice (or *Disc1* heterozygous mice) to mimic patients with psychiatric disorders who exhibit decreased GLO1 activity or expression (or decreased *Disc1* expression), whereas high-sucrose intake was used as an environmental risk factor (bottom panel). We investigated four groups of mice: WT, starch-fed mice (Starch +/+); *Glo1* (or *Disc1*) heterozygous, starch-fed mice (Starch *Glo1*/+ (or Starch *Disc1*/+)); WT, sucrose-fed mice (Suc +/+); and *Glo1* (or *Disc1*) heterozygous, sucrose-fed mice (Suc *Glo1*/+ (or Suc *Disc1*/+)).

### Animals

All experimental procedures were approved by the Animal Experimentation Ethics Committee of the Tokyo Metropolitan Institute of Medical Science (49040). All mice were maintained under a 12:12 h light:dark cycle (lights on at 8:00 AM) with free access to the indicated diet. All efforts were made to minimize the number of animals used and their suffering. *Glo1* knockout mice were generated as described previously (*54, 55*). Briefly, *Glo1*-trapped ES cell lines from the International Gene Trap Consortium were used for the generation of 3 founder mice, which were then backcrossed to C57BL/6 mice. In the manuscript, trapped *Glo1* is referred to as *Glo1*. *Disc1-LI* (locus-impairment) heterozygous mutant mice were generated as previously described (*16*). In the manuscript, *Disc1-LI* is referred to as *Disc1*. Alternatively, mice were backcrossed to GFAP-GFP mice to monitor astrocyte activation (*31, 32*). Male mice were exclusively used in the behavioral tests, whereas mice of both sexes were used in histological, biochemical, and physiological experiments.

### Diet preparation

The two diets used in the present study were newly created in collaboration with Oriental Yeast Co. Ltd. (Tokyo, Japan). We named the sucrose diet HSD-70 (# OYC 2405100) and the starch diet HCD-70 (# OYC 2405000) (Supplementary Table 4). They contained the same caloric proportions of carbohydrate, fat, and protein; however, all carbohydrate calories are derived from either starch or sucrose.

### Drug preparation

Aripiprazole was dissolved in acetic acid and diluted to 3.5 mg/L in water for administration at 0.5 mg/kg/day. The final acetic acid concentration in the drinking water was 0.7%. Aspirin was dissolved in ethanol and diluted to 70 mg/L in water for administration at 1 mg/kg/day. The final ethanol concentration in the drinking water was 0.15%. The daily dose was based on the measurement of mean water consumption daily in all sucrose-fed *Glo1* heterozygous mice and starch-fed wild-type (WT) mice (fig. S1D, E). There were no between-group differences in terms of aripiprazole consumption (Student’s *t*-test, Starch +/+ and Suc *Glo1*/+: *p* = 0.86) or aspirin-containing water (Student’s *t*-test, Starch +/+ and Suc *Glo1*/+: *p* = 0.61).

### Behavioral tests

Mice were habituated in the behavioral room for >30 min before each test. Behavioral tests were performed in the following sequence of increasing stress: elevated plus maze, grooming, nest building, open field, object location, social interaction, and pre-pulse inhibition (PPI). All test apparatuses were cleaned using 70% ethanol and water between trials, and the subsequent test session was started only after the ethanol vapor odors had disappeared and the apparatuses had dried.

The elevated plus maze (EPM-04M, Muromachi, Japan) consisted of two opposing open arms (297 × 54 mm) and two closed arms (300 × 60 × 150 mm) extending from a central platform (60 × 60 mm). The entire apparatus was elevated 400 mm above the floor. Each mouse was placed on the central platform facing a closed arm and allowed to freely explore the maze for 10 min. Arm entry was defined as the entry of all four paws into the arm. The time spent in the open arms over 10 min was recorded as an index of state anxiety.

For the self-grooming test, all mice housed in the same home cage were moved into a new cage for 10 min. Each mouse was then individually placed in a standard mouse home cage (31 × 16.5 × 14 cm) illuminated at ∼200 lux. After a 10-min habituation period, each mouse was scored for cumulative time spent grooming all body regions (*56*) over 10 min using a stopwatch. Self-grooming behavior is conserved across species and is indicative of certain pathological conditions or factors. In humans, for example, self-grooming increases during stressful conditions and in certain psychiatric disorders (*56*).

For the nest building test, 200 g of corncob was spread across the bottom of each cage for bedding, and a square-shaped piece of cotton was placed in the cage center as raw material for the nest. Each mouse was individually placed in the cage for 8 h. Photos of the constructed nest were acquired every 2 h, and the nest building process was evaluated by measuring the proportion of loose cotton as follows: 1 point for 25% weight (Wt%) loosened, 2 points for 50 Wt% loosened, 3 points for 75 Wt% loosened, and 4 points for 100 Wt% loosened. After 8 h, we checked the shape of the nest and added 1 point if the mice had completed a nest with a bird’s nest-like shape. The temperature of the room was maintained at 25°C and illumination at 150–180 lux during nest building. Nest-building behavior is an indicator of general well-being in mice (*57*), whereas poor nest building is an indicator of psychological or physiological abnormalities (*58, 59*).

For the open field (OF) test, each mouse was placed in the center of the apparatus (40 × 40 × 40 cm; 150–180 lux illumination) and allowed to move freely for 10 min. The behavior of each mouse was monitored using a charge-coupled device camera mounted on the ceiling above the OF. The total distance traveled (cm) was measured using CompACT VAS software (Muromachi).

For the object location test (OLT) of working memory (*60*), mice first explored the empty OF box, and then, two identical objects A and B (two 500-mL PET bottles filled with blue-colored water) were placed in two corners 5 cm from the wall. After a 10-min exploration/learning period, the mice were returned to their home cage for 5 min, and Object A was moved to a new corner (Object A). The animals were then placed back in the ′ OF box and allowed to explore for 5 min. The time spent exploring A and B were ′ measured to calculate a discrimination index representing working memory according to the following equation: Discrimination Index = (Novel Object A exploration time Familiar Object B exploration time) / (Novel Object A + Familiar Object B exploration times). The ′ OLT was performed under illumination at 10–15 lux.

The social interaction test was conducted as previously described (*61*) using a specialized Sociability Apparatus (SC-03M, Muromachi). The time spent sniffing a novel stimulus mouse or object was manually scored from videos recorded using an overhead color USB camera (aa000080a02, Iroiro House). Stimulus mice (129Sv/SvJcl strain) age- and sex-matched to test mice were habituated to the apparatus and to the enclosure cup for 30 min per day for 2 days prior to testing. The location (left or right) of the novel object and novel mouse within an enclosure were alternated across test subjects. The test mouse was allowed to acclimate to the apparatus for a total of 20 min before the sociability test—the first 10 min in the central chamber with the doors closed and then 10 min in the empty arena with the doors open. The test subject was briefly confined to the center chamber while a novel stimulus mouse in an enclosure cup was placed on one of the side chamber and another empty enclosure cup (novel object) was placed on the other side of the chamber. The test subject was allowed to approach the novel object or mouse freely for 10 min. The time spent interacting with the stimulus mouse versus the novel object was calculated as an index of sociability.

The SR-LAB-Startle Response System (San Diego Instruments) was used to detect acoustic startle reflexes and PPI. Startle responses were measured using 5 stimulus intensities (80, 90, 100, 110, and 120 dB) delivered 10 times each for 40 ms over a white noise background (65 dB). The stimuli were presented in quasi-random order at random inter-trial intervals (10–20 s). In the PPI session, mice were exposed to 2 stimulus patterns: 1) a startle stimulus alone (120 dB, 40 ms) with no pre-pulse stimulus and 2) a startle stimulus (120 dB, 40 ms) following a pre-pulse stimulus (70 dB for 20 ms; lead time, 100 ms). Each trial was repeated 10 times in quasi-random order at random inter-trial intervals (10–20 s). PPI was defined as the percent decline in startle response because of pre-pulse stimuli according to the following equation: 100 − [(120 dB startle amplitude after any pre-pulse) / (120 dB startle amplitude only)] × 100. A new accelerometer was used for the experiments with *Disc1* knockout mice.

### Immunohistochemistry

Following transcardial perfusion with PBS and 4% paraformaldehyde, entire brains were collected, post-fixed at 4°C overnight, and cryoprotected in 20% sucrose at 4°C overnight. Serial coronal sections (50 μm) were then cut using a cryostat (CM3050 S; Leica Microsystems). The antigens in the tissues were reactivated by heating in HistoVT One solution (Nacalai Tesque) for 30 min at 70°C using a water bath. Sections were permeabilized with 0.2% Triton X-100 and 1% Block Ace (DS Pharma Biomedical) in PBS for 30 min at room temperature, following which they were incubated overnight with the indicated primary antibodies at room temperature. For immunohistochemistry of postmortem human brain tissues, paraffin blocks including BA9 (a region of frontal cortex) were sliced into 10-μm sections, deparaffinized with xylene, and rehydrated with decreasing concentrations of ethanol in water. Antigens were reactivated by heating in HistoVT One solution for 30 min at 90°C using a water bath. Sections were treated with TrueBlack Lipofuscin Autofluorescence Quencher (Biotium Inc.) for 30 s at room temperature and blocked with 1% Block Ace (DS Pharma Biomedical) in PBS for 30 min at room temperature. Thereafter, the mouse and human brain sections were subjected to the same immunostaining procedures. The following primary antibodies were diluted in PBS containing 0.4% Block Ace: goat anti-PV (Frontier Institute, PV-Go-Af860; 1:2000), mouse anti-ALDH1L1 (Rockland, 600-101-HB6S; 1:200), FITC-conjugated tomato lectin (VECTOR, FL-1171; 1:200), chick anti-GFP (Abcam, ab13970; 1:500), goat anti-IBA1 (Abcam, ab48004; 1:100), mouse anti-NeuN (Millipore, MAB377; 1:500), rabbit anti-AGE (Abcam, ab23722; 1:2000), rabbit anti-IBA1 (Wako, WDJ3047; 1:300), rat anti-CD68 (Abcam, ab53444; 1:500), and rabbit anti-fibrin (Dako, A0080, 1:500). Thereafter, sections were washed three times with PBS containing 0.05% Tween-20, incubated for 2 h with fluorochrome-conjugated secondary antibodies in PBS containing 0.4% Block Ace, and washed an additional three times in PBS containing 0.4% Block Ace. For enhanced horseradish peroxidase (HRP) immunostaining, samples were treated with 3% H_2_O_2_ in PBS for 20 min following the reactivation step to quench endogenous peroxidase activity and washed in PBS. Sections were incubated with rabbit anti-GLO1 (Novusbio, NBP2-75514, 1:1500) and/or mouse anti-AGE4 (Trans Genic Inc, 14B5, 1:400), followed by incubation with anti-IgG antibodies conjugated to biotin (Vector, 1:200). After washing as described for other secondary antibodies, sections were incubated with streptavidin-conjugated HRP (Jackson ImmunoResearch, 1:200) for 120 min and washed three times with PBS containing 0.05% Tween-20. The TSA Plus Fluorescence System (PerkinElmer) was used to detect HRP activity. All preparations were counterstained with DAPI (Nacalai Tesque) to reveal cell nuclei, washed three additional times, mounted in Permaflow (Thermo Scientific), and observed using a FluoView^®^ FV3000 Confocal Laser Scanning Microscope (Olympus).

### Image analysis

Unless otherwise noted, all image analyses were performed using ImageJ version 2.0.0-rc-59/1.51n, and images were first binarized. PV-positive cells with a threshold exceeding 30 were counted.

To measure the areas immunopositive for AGE, fibrin, and AGE-4, the threshold settings were applied to ensure that only the correct immunopositive area was properly selected, and the stained area of the entire screen was measured. For measuring the fluorescence intensity of GFP and IBA1, after setting the ROI to select only target immunoreactive cells, the intensity of >5 cells in each image were measured. To count microglial cells containing a CD68-positive area exceeding 20 µm^2^, the threshold was set to select only the CD68-positive area. Colocalization analysis of two different fluorescence markers was performed using the Fiji plug-in Coloc 2 with the default settings. We adopted Pearson’s R value below the threshold to judge the strength of colocalization: R ≥ 0.7, strong; 0.7 > R ≥ 0.4, moderate; 0.4 > R ≥ 0.2, weak; and R < 0.2, none or very weak.

### Immunoblotting

Extracts from mouse hippocampi were homogenized in lysis buffer containing 40 mM Tris base, 0.4% sodium dodecyl sulfate (SDS), 0.01 M EDTA (pH 8.0), 8 M urea, and 1 mM phenylmethylsulfonyl fluoride. The total lysate protein content was quantified using a DC Protein Assay Kit (Bio-Rad). Total protein (30 µg per gel lane) was separated using SDS-PAGE and transferred to PVDF membranes (Millipore). Membranes were blocked with TBST buffer (137m M NaCl, 2.7 mM KCl, and 25mM Tris, pH 8.0) including 0.2% Triton X-100 and 5% bovine serum albumin (BSA) for 30 min at room temperature with slow shaking, followed by incubation overnight with primary antibodies in TBST including 2% BSA at 4°C. The primary antibodies used were rabbit anti-GLO1 (Santa Cruz, sc-67351; 1:1000), mouse-anti-PV (Swant, PV-235; 1:1000), rabbit anti-glucose transporter 1 (Glut1, Frontier Institute, Af1020; 1:1000) and mouse anti-tubulin (Santa Cruz, sc-32293; 1:10000). After washing three times with TBST, membranes were incubated with the secondary antibody (HRP-conjugated anti-mouse or anti-rabbit IgG antibody, GE Healthcare; 1:2000) in TBST plus 2% BSA. After washing three times with TBST, blots were processed for chemiluminescence using standard protocols (ECL Prime Western Blotting Detection Regent #RPN2236, GE Healthcare), and signals were detected using a LAS 4000 Imager (Fuji Film).

### Microdialysis

We used an *in vivo* microdialysis system for the measurement of extracellular dopamine concentration (*62*) and collection of brain parenchymal dialysate (Fig.1H, J and fig. S7D). After anesthesia by intraperitoneal injection of ketamine (80 mg/kg)/xylazine (16 mg/kg), mice were fixed in a stereotaxic apparatus (Narishige) and a microdialysis guide cannula (CXG-8, Eicom) was implanted in the medial prefrontal cortex (mPFC; antero-posterior (AP), +1.8 mm; medio-lateral (ML), ±0.15 mm; dorso-ventral (DV), −1.5 mm from bregma), or nucleus accumbens (NAc) (AP, +1.5 mm; ML, ±0.6 mm; DV, −3.5 mm from bregma). After recovery for at least 10 days, a microdialysis probe (CX-I-8-01 for the mPFC and CX-I-8-02 for NAc; Eicom) was inserted via the guide cannula. Following insertion, the probe was connected to a syringe pump and perfusion was performed at 2 μ L/min for NAc and 0.5 μL/min for mPFC using Ringer’s solution (147 mM NaCl, 4 mM KCl, and 2.3 mM CaCl_2_). Dialysate samples were collected every 10 min and automatically loaded onto an HTEC-500EPS HPLC unit (Eicom). Constant 5-HT concentration in three consecutive collection periods was first confirmed to rule out blood contamination before initiating the measurements of dopamine concentration or collection of parenchymal dialysates. Analytes were then separated on an affinity column (PP-ODS III, Eicom), and compounds were subjected to redox reactions within an electrochemical detection unit (amperometric DC mode; applied potential range, 450 mV). The resulting chromatograms were analyzed using an EPC-500 data processor (Eicom), and actual sample concentrations were computed based on the peak heights obtained using 0.01, 0.1, and 1 pg dopamine in standard solution (Sigma). The locations of the microdialysis probes were histologically confirmed.

For glucose measurements in Fig. 4H and Fig. 5G, we collected dialysates from the mPFC for 1 h after 16 h of fasting, followed by 0–1 h after eating 0.05 g of carbohydrates (starch or sucrose), and 1–2 h after eating 0.05 g of carbohydrates (starch or sucrose).

### EEG recordings

For behavioral and video/EEG monitoring, mice were anesthetized by an intraperitoneal injection of ketamine (80 mg/kg)/xylazine (16 mg/kg), fixed in a stereotaxic apparatus (Narishige, Japan), and implanted with EEG and electromyography (EMG) electrodes. The EEG electrodes were gold-coated stainless steel screws (SUS303) soldered with lead wires (ANE-0190, Adler’s Nest, Japan) implanted epidurally over the left frontal cortex (AP, 1 mm; ML, 1 mm) and the bilateral parietal cortex (AP, −2 mm; ML, ±2 mm). All wires were soldered to a multichannel electrical connector (R833-83-006-001, TOKIWA SHOKO, Japan). The left parietal cortex electrode was used as a reference for monitoring the frontal cortex EEG. The EMG electrodes were lead wires placed bilaterally into the trapezius muscle. Following recovery for at least 10 days, EEG/EMG signals were amplified and band-pass filtered (EEG: 1.5–1000 Hz; EGM: 15–3000 Hz) using a MEG-6116 system (NIHON KOHDEN), digitized at a sampling rate of 200 Hz, recorded using a data acquisition system (PowerLab 8/30, ADInstruments), and analyzed using LabChart Software (ADInstruments). Behavioral activities were recorded using a USB camera (aa000080a02, Iroiro House, Japan). Behavioral and electrophysiological responses to a novel object (an empty 100 mL DURAN bin) were recorded in an OF chamber (20 × 20 × 26 cm). The novel object was placed in one corner of the OF chamber to induce exploration. The 30 s preceding the first contact with the novel object was analyzed for object recognition (“object activity”). For EEG monitoring in the home cage, mice were first habituated for 8 h. Home cage EEG data were then acquired for 2 min after awaking as confirmed by clear EMG signals and movement images from an offline video camera analysis (“home cage activity”). All recordings were converted into power spectra using a fast Fourier transform (FFT) algorithm with a 5-s Hann cosine-bell window and 50% overlap between successive window measurements. All FFTs were maintained at 1024 points to obtain 0.512 Hz resolution. The total signal amplitude or power (V^2^) in each 5-s period was measured as the power magnitude at each frequency. The mean of the grouped power spectra was calculated over the following frequency ranges: 1–4 Hz (delta), 5–10 Hz (theta), 30–45 Hz (low gamma), and 55–80 Hz (high gamma). The power values detected at each frequency range for 30 s were further averaged over 30 s of total EEG power using the mean values to remove potential noise. These analyses were performed using custom software written in MATLAB (R2019b; MathWorks).

### Transcriptome analysis

Three independent total RNA samples from each group were mixed and purified using a RNeasy Mini Kit (Qiagen). RNA quality was assessed using a 2100 bioanalyzer (Agilent Technologies). Cy3-labeled cRNA was prepared using a Low Input Quick Amp Labeling Kit (Agilent Technologies), in accordance with the manufacturer’s protocol. Samples were hybridized to the SurePrint G3 Mouse Gene Expression v2 Microarray (G4852B; Agilent Technologies). Thereafter, the array was washed and scanned using the SureScan Microarray Scanner (Agilent Technologies). Microarray images were analyzed using the Feature Extraction software with default settings for all parameters (Agilent Technologies). Data from each microarray analysis were normalized by shift to the 75^th^ percentile without baseline transformation. Microarray results were deposited in the Gene Expression Omnibus database under the accession number GSE141829.

### Insulin and glucose measurements

Blood plasma was collected from the mouse cheek as described by Golde (*63*). Plasma glucose concentration was measured using a Precision-Neo blood glucose meter (#71386-80, Abbott Japan), plasma insulin concentration using an ELISA kit (#M1102, MORINAGA), and glucose concentration in the dialysate samples using a different ELISA kit (#ab65333, Abcam), all according to the manufacturers’ instructions. Data were collected on a microplate reader (Varioskan, Thermo Fisher Scientific).

### Human postmortem brain tissue collection

Postmortem brain tissues from patients with SZ and BD were obtained from the Fukushima Brain Bank at the Department of Neuropsychiatry, Fukushima Medical University. Postmortem brain tissues from control individuals were obtained from the Section of Pathology, Fukushima Medical University Hospital. The use of postmortem human brain tissues in the present study was approved by the Ethics Committee of Fukushima Medical University (No.1685) and Tokyo Metropolitan Institute of Medical Science (No. 18-20) and complied with the Declaration of Helsinki and its later amendments. All procedures were conducted with the informed written consent of the next of kin. Detailed demographic information of the 10 patients with SZ, 9 patients with BD, and the 12 age- and sex-matched control individuals is provided in Supplementary Table 3. No between-group differences were observed in terms of sex (Fisher’s exact test, Ctrl and SZ: *p* = 1.00, Ctrl and BD: *p* = 0.40), age (Student’s t-test, Ctrl and SZ: *p* = 0.69, Welch’s t-test, Ctrl and BD: *p*= 0.66), postmortem interval (Student’s t-test, Ctrl and SZ: *p* = 0.89, Ctrl and BD: *p* = 0.98), or a history of diabetes mellitus (Fisher’s exact test, Ctrl and SZ: *p* = 0.59, Ctrl and BD: *p* = 0.59). Each patient with SZ and BD fulfilled the diagnostic criteria established by the American Psychiatric Association (Diagnostic and Statistical Manual of Mental Disorders, DSM-IV) and did not have a past history of other neurological disorders or substance abuse. Moreover, none of the control individuals had any record of mental disorders, neurological disorders, or substance abuse.

### ROS detection

DHE is oxidized by superoxide and other reactive species to produce a precipitate that emits red fluorescence, whose intensity can be used to measure the degree of cellular oxidative stress. Animals were quickly perfused with ice-cold phosphate-buffered salts (PBS), and the decapitated brains were embedded in OCT compound (Sakura Finetek Japan), and 10 μm thick coronary sections were prepared in a cryostat. These sections were incubated with phosphate buffer containing 100 μM diethylene triamine pentaacetic acid (PB/DTPA) for 10min, and then incubated with PB/DTPA containing 10 μm DHE (Invitrogen) for 20 min. After washing with PB/DTPA twice, sections were fixed with 4% PFA for 15 min. Once washed with PBS, PBS containing 1 µM TO-PRO 3 (Thermo Scientific) was applied to each section followed by 3 washes with PBS. Then, sections were sealed with Permaflow (Thermo Scientific) and observed using a FluoView® FV3000 Confocal Laser Scanning Microscope (Olympus). For DHE detection, we used the following filter set: excitation 405 nm, emission 600–650 nm.

### Statistical analyses

Statistical differences among ≥4 groups were determined using one-way analysis of variance (ANOVA), two-way ANOVA, three-way ANOVA, or repeated measures ANOVA, followed by the Bonferroni multiple comparison test or Tukey-Kramer test as a post hoc test, as summarized in Supplementary Table 5. We used the same animals for all results coming from Starch +/+ and Suc *Glo1*/+ groups according to the 3R rule. Data were collected at the same time regardless of whether the drug was administered or not. The number of animals employed for each experiment/analysis can be found in each figure legend. Detailed descriptions of statistical analyses in Fig.1H, J, and 4H are shown below. In Figure 1H, post hoc Dunnett‘s multiple comparisons test for groups was performed at specific time points (0 min and 40 min), ^###^*p* < 0.001 for Suc *Glo1*/+ vs. Ctrl (Starch +/+), ^$^*p* < 0.05 for Starch *Glo1*/+ vs. Ctrl (Starch +/+). In Fig. 1J, post hoc Bonferroni’s multiple comparisons test for groups was performed at specific time points (0 min and 40 min), ^###^*p* < 0.001 for Suc *Glo1*/+ vs. Starch +/+, ^$$$^*p* < 0.001 for Suc *Glo1*/+ vs. Starch +/+ Aripi, ^&&&^*p* < 0.001 for Suc *Glo1*/+ vs. Suc *Glo1*/+ Aripi. In Fig. 4H, post hoc Dunnett’s multiple comparisons test for groups was performed at each time point, ^###^*p* < 0.001, ^##^*p* < 0.01 for Suc *Glo1*/+ vs. Ctrl (Starch +/+).

## Supplementary Figure Legends

**Supplementary Figure 1.**
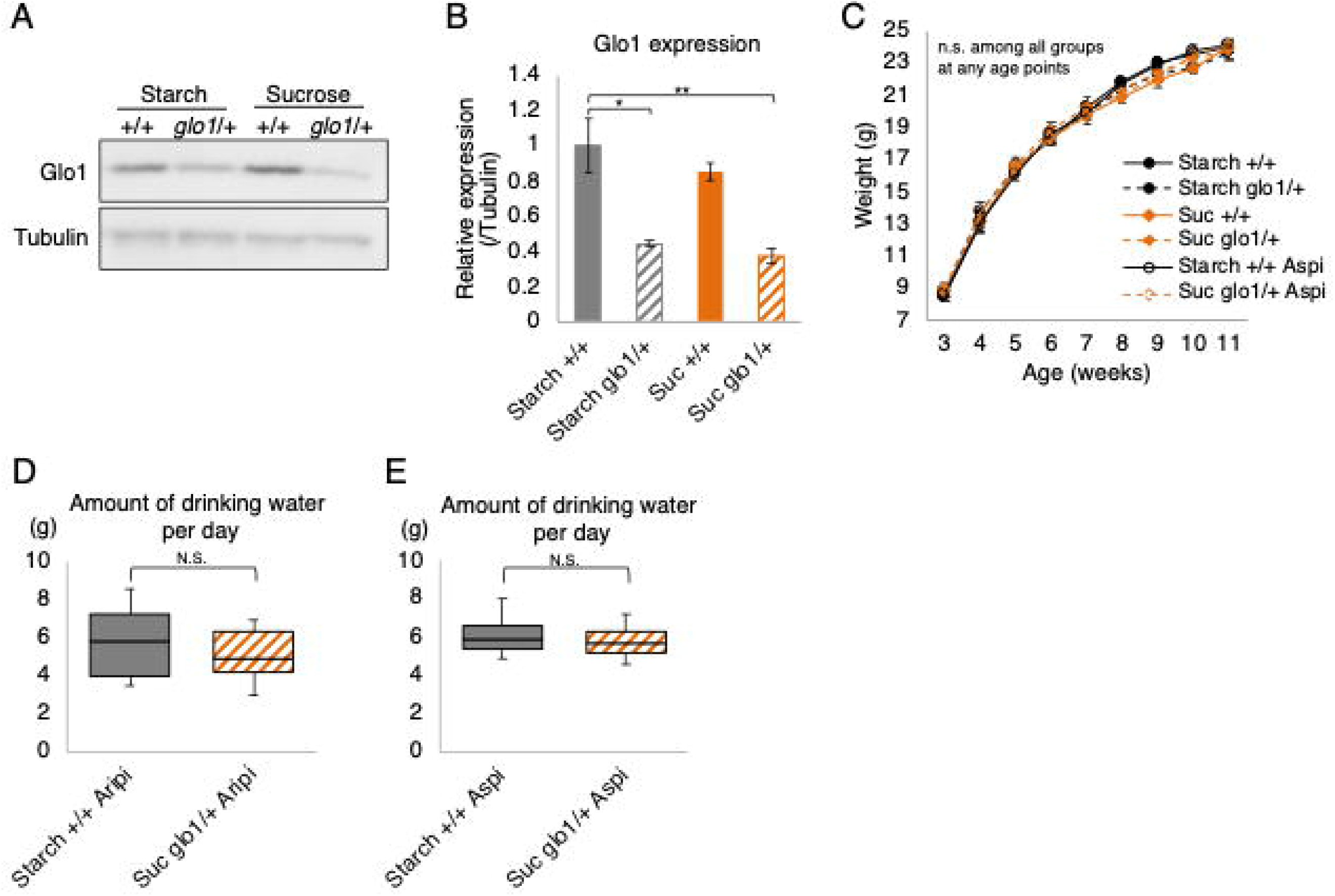
Characterization of regional GLO1 expression in WT and heterozygous *Glo1* mutant mice fed starch or sucrose. (**A**) Western blot analysis of GLO1 protein expression using tubulin as internal control. The cerebral cortex, including the hippocampus, was used as loading sample. (**B**) Densitometric analysis of GLO1 protein expression (n = 3 or 4 mice per group). To quantify expression, GLO1 band intensities in (**A**) were divided by the intensities of the corresponding tubulin bands. (**C**) Body weight trajectories. No significant differences were observed among groups (n = 6–10 mice per group). (**D, E**) Measurement of aripiprazole- or aspirin-containing water consumption per day to estimate drug intake (n = 4 mice per group). Student’s *t*-test was used for statistical analysis. All data are presented as mean ± SEM. \*\**p* < 0.01, \**p* < 0.05

**Supplementary Figure 2.**
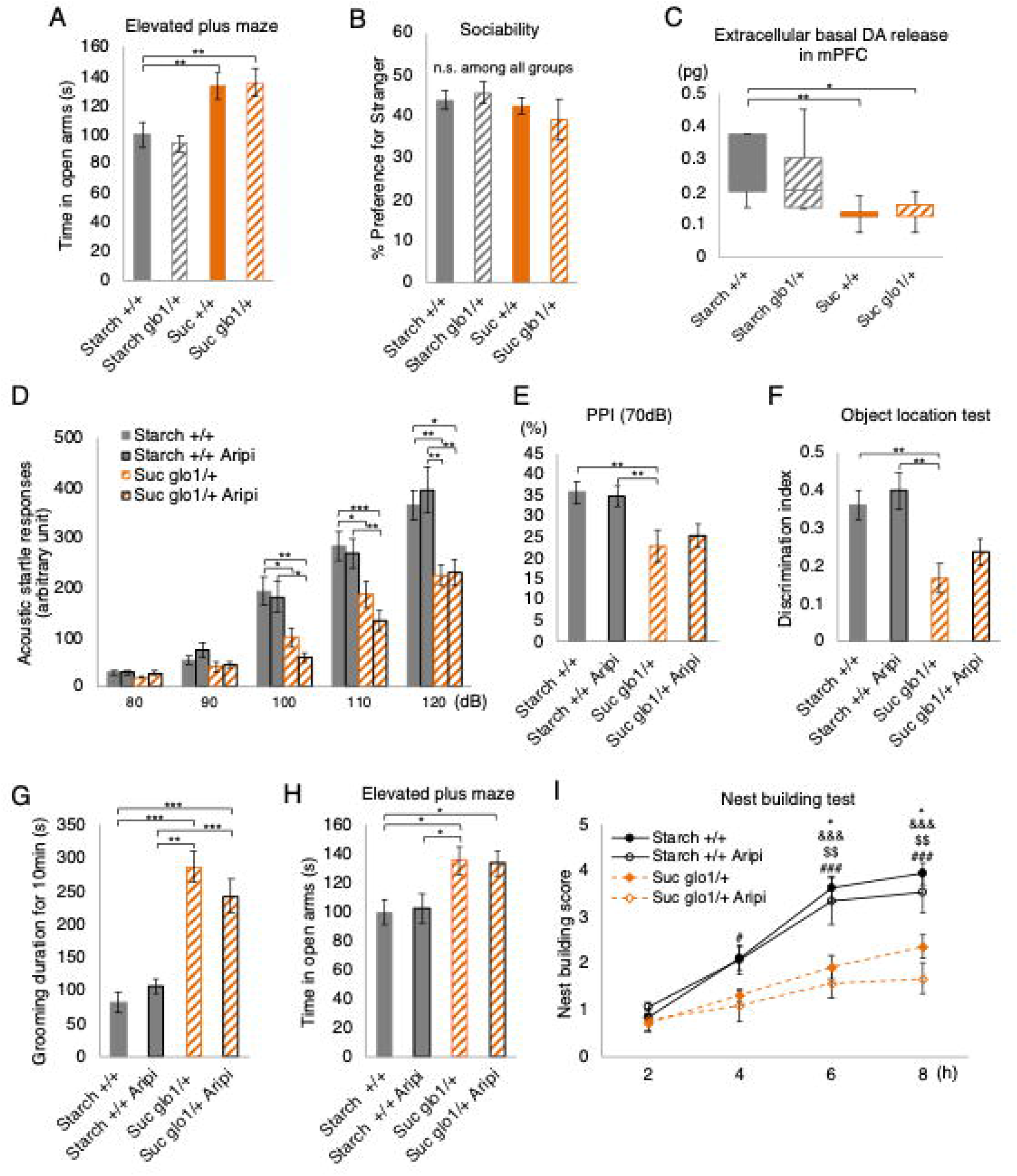
Analyses of psychiatric disease-related phenotypes in *Glo1* heterozygotic mice. (**A, B**) Behavioral analyses in the four groups of mice (n = 18–23 mice per group). (**A**) Elevated plus maze test to evaluate anxiety. (**B**) Preference for novel mouse in the three-chamber test, calculated as [(time spent exploring novel mouse)/(total time spent exploring novel mouse and novel object)]□× □100%. (**C**) Extracellular dopamine concentration in the medial prefrontal cortex measured at 20-min intervals using an *in vivo* microdialysis system (n = 7–10 mice per group). (**D–I**) Effect of aripiprazole treatment on behavioral phenotypes (n = 12–18 mice per group). (**D**) Effects of aripiprazole treatment on the acoustic startle response, (**E**) pre-pulse inhibition (PPI) using a 70-dB pre-pulse, (**F**) object location test, (**G**) self-grooming, (**H**) elevated plus maze performance, and (**I**) nest-building skill. In **(I)**, post hoc Tukey‘s multiple comparisons test of groups at specific time points, ^###^*p* < 0.001, ^#^*p* < 0.05 for Starch +/+ vs. Suc *Glo1*/+ Aripi, ^$$^*p* < 0.01 for Starch +/+ Aripi vs. Suc *Glo1*/+ Aripi, ^$$$^*p* < 0.001 for Starch +/+ vs. Suc *Glo1*/+, \**p* < 0.05 for Starch +/+ Aripi vs. Suc *Glo1*/+. All data are presented as mean ± SEM. \*\*\**p* < 0.001, \*\**p* < 0.01, \**p* < 0.05

**Supplementary Figure 3.**
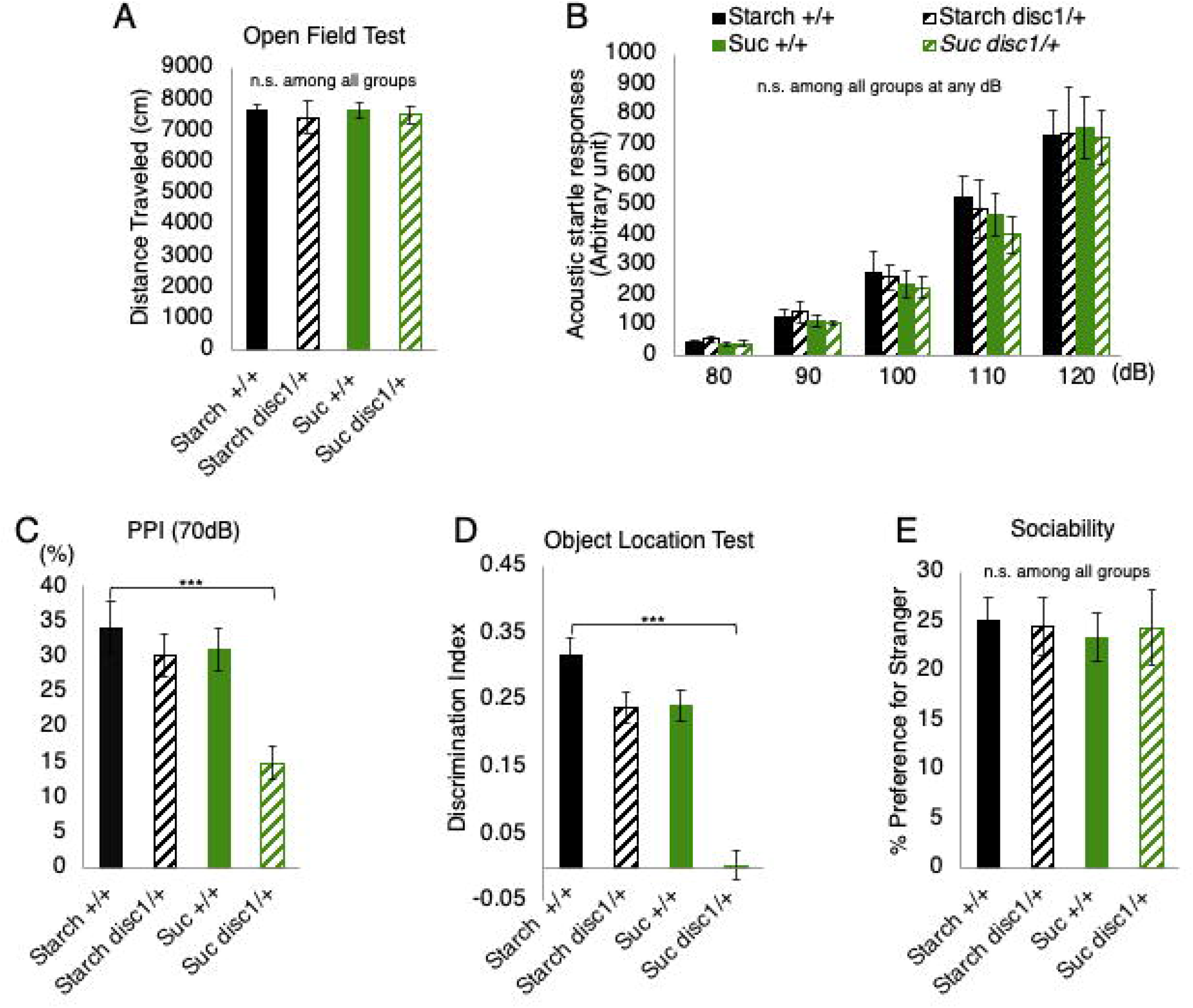
Analyses of psychiatric disease-related phenotypes in *Disc1* heterozygotic mice. (**A-E**) Behavioral analyses in the four groups of mice (n = 8–11 mice per group). (**A**) Spontaneous locomotor activity in the open-field test. (**B**) Acoustic startle responses. (**C**) Pre-pulse inhibition (PPI) using a 70 dB pre-pulse. (**D**) Object location test. (**E**) Preference for novel mouse in the three-chamber test, calculated as fig. S2B. All data are presented as mean ± SEM. \*\*\**p* < 0.001

**Supplementary Figure 4.**
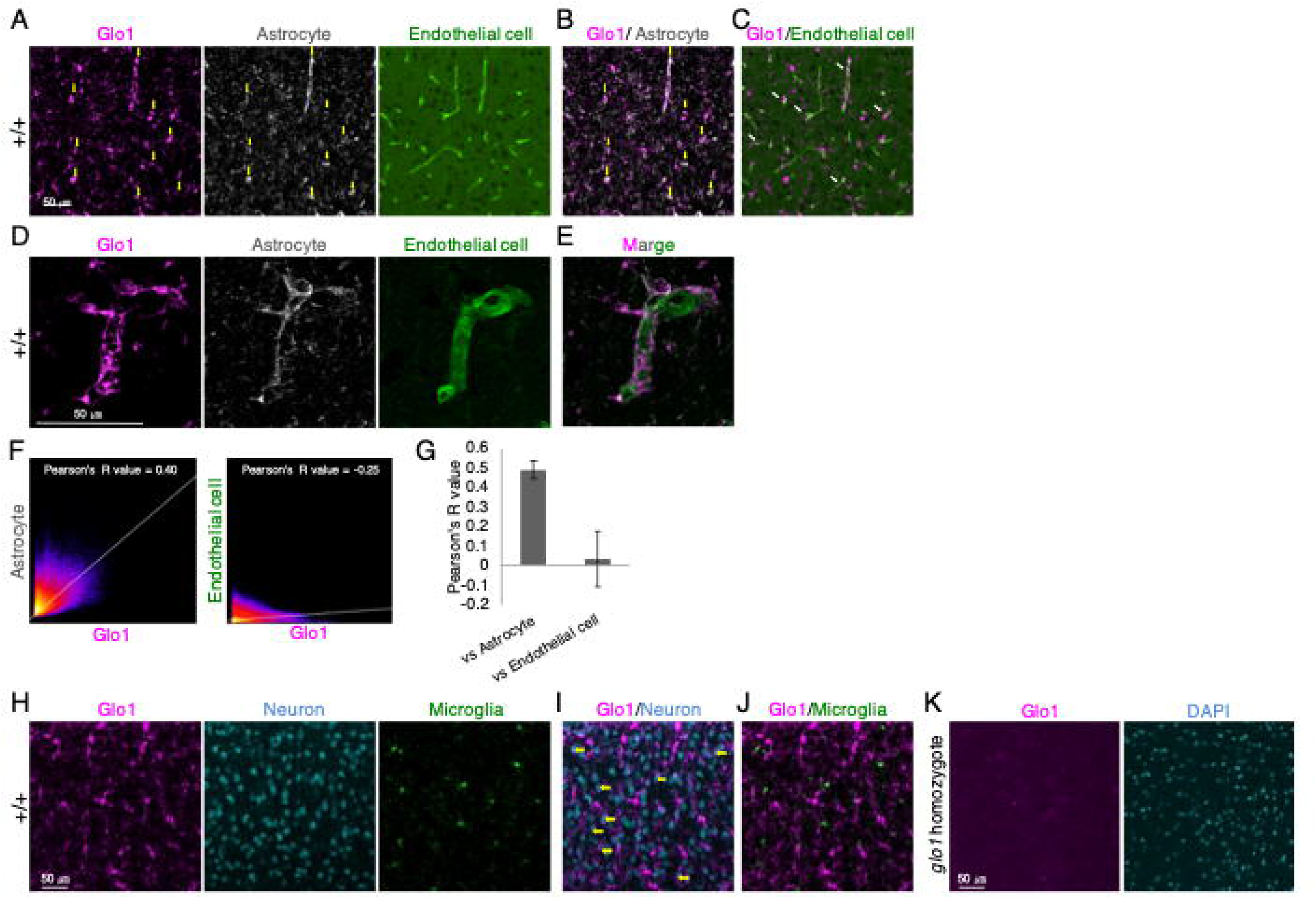
GLO1 cellular localization in the prefrontal cortex of sucrose-fed wild-type mice. (**A–F**) GLO1 localization in astrocytes. (**A**) GLO1 co-immunostaining with an astrocyte marker (ALDH1L1) or an endothelial cell marker (tomato lectin). (**B**) Merged GLO1/ALDH1L1 image from (**A**). Yellow arrows in (**B**) indicate cells with GLO1/ALDH1L1 colocalization. (**C**) Merged GLO1/lectin image from (**A**). White arrows denote representative GLO1-positive cells close to endothelial cells. (**D, E**) High magnification images of GLO1 co-immunostaining with ALDH1L1 or with tomato lectin. (**E**) Merged GLO1/ALDH1L1 image from (**D**). (**F**) 2D intensity histogram of the two indicated channels to identify GLO1 colocalization with ALDH1L1 or tomato lectin, interpreting the strength of a relationship based on R as follows: R ≥ 0.7, strong; 0.7 > R ≥, moderate; 0.4 > R ≥ 0.2, weak; and R < 0.2, none or very weak. (**G**) Average R-value of colocalization data including (**F**) from three different locations of three mice. (**H**) GLO1 co-immunostaining with the neuronal marker NeuN and the microglial marker IBA1. (**I**) Merged GLO1/NeuN image from (**H**). Yellow arrows indicate neurons with mild GLO1 immunoreactivity. (**J**) Merged GLO1/IBA1 image from (**H**). (**K**) GLO1 immunostaining and DAPI staining in *Glo1* homozygous mice.

**Supplementary Figure 5.**
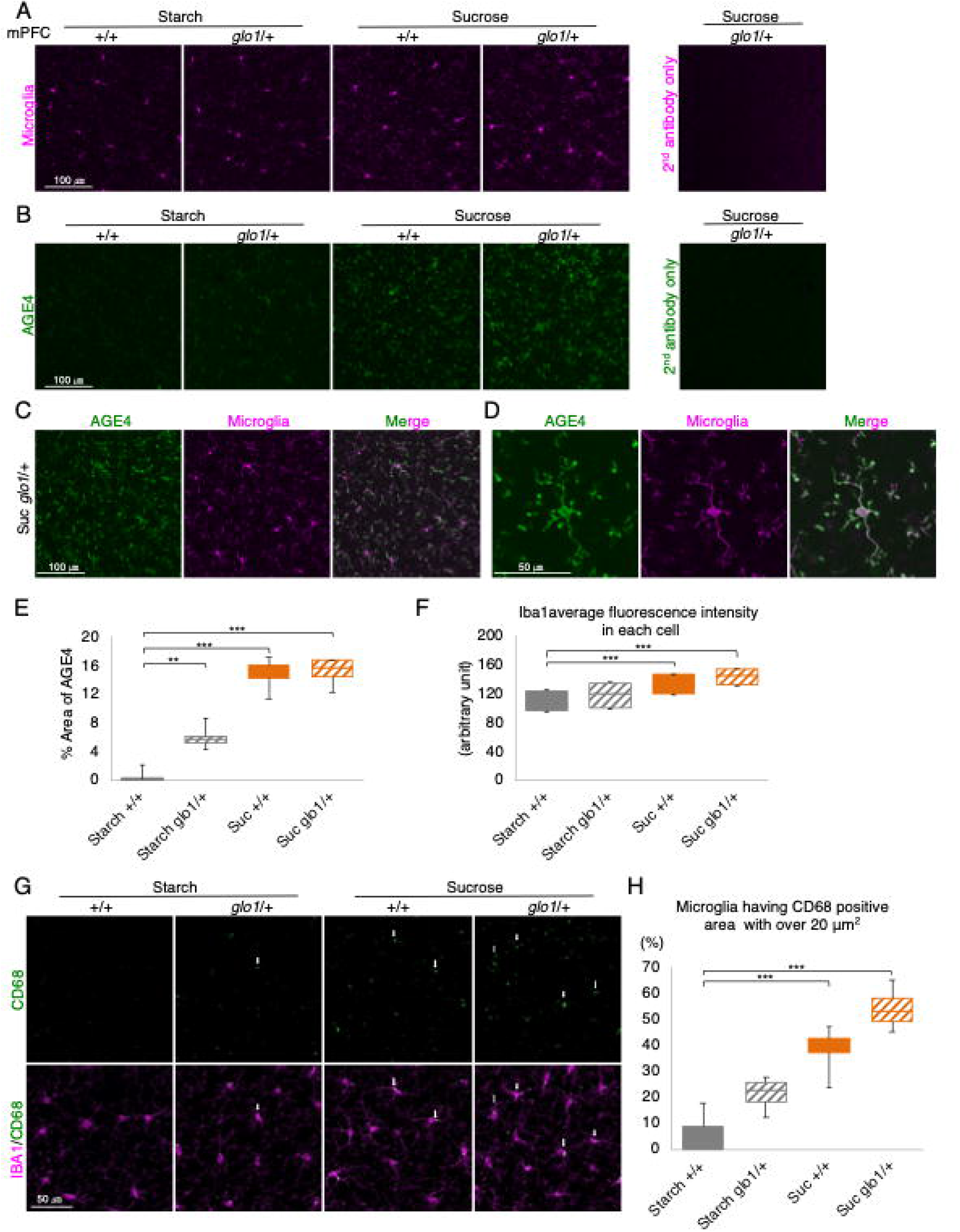
Fructose-derived AGE4 accumulation in microglia of sucrose-fed mice. (**A, B**) Immunohistochemical images of the microglial marker IBA1 and AGE4 in the medial prefrontal cortex region (mPFC). (**C**) Merged AGE4/IBA1 image. (**D**) High magnification images of AGE4 co-immunostaining with IBA1. (**E**) Measurement of the area of AGE-4 immunoexpression in (**B**). The mean AGE4-immunopositive area intensity of the entire image was measured in each section (n = 4 mice per group). (**F**) Mean IBA1 fluorescence intensities of five randomly selected cells per image in four independent mice. (**G**) Immunohistological images of IBA1- and CD68-positive microglia in the mPFC region. White arrows in (**G**) indicate microglial cells containing CD68-positive areas >20 µm^2^. (**H**) Percentage of CD68-positive cells per total microglial cells in each image in (**G**). All data are presented as mean ± SEM. \*\*\**p* < 0.001, \*\**p* < 0.01

**Supplementary Figure 6.**
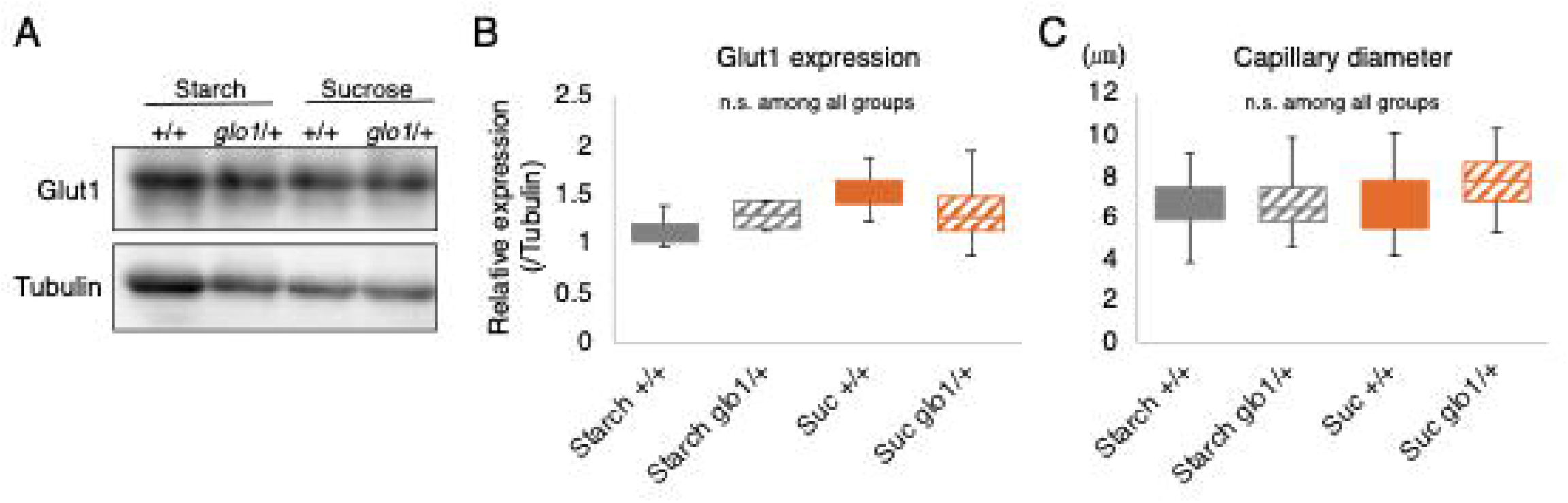
Blood vessels Characterization in the four groups. (**A**) Western blot analysis of glucose transporter 1 (Glut1) protein expression using tubulin as internal control. The cerebral cortex, including the hippocampus, was used as loading sample. (**B**) Densitometric analysis of Glut1 protein expression (n = 3 mice per group). To quantify expression, Glut1 band intensities in (**A**) were divided by the intensities of the corresponding tubulin bands. (**C**) Measurement of capillary diameter. Vessels <10 µm in diameter were defined as capillaries, and the mean diameter of 10 randomly selected capillaries per image measured in Fig. 4C (n = 3 mice per group). All data are presented as mean ± SEM.

**Supplementary Figure 7.**
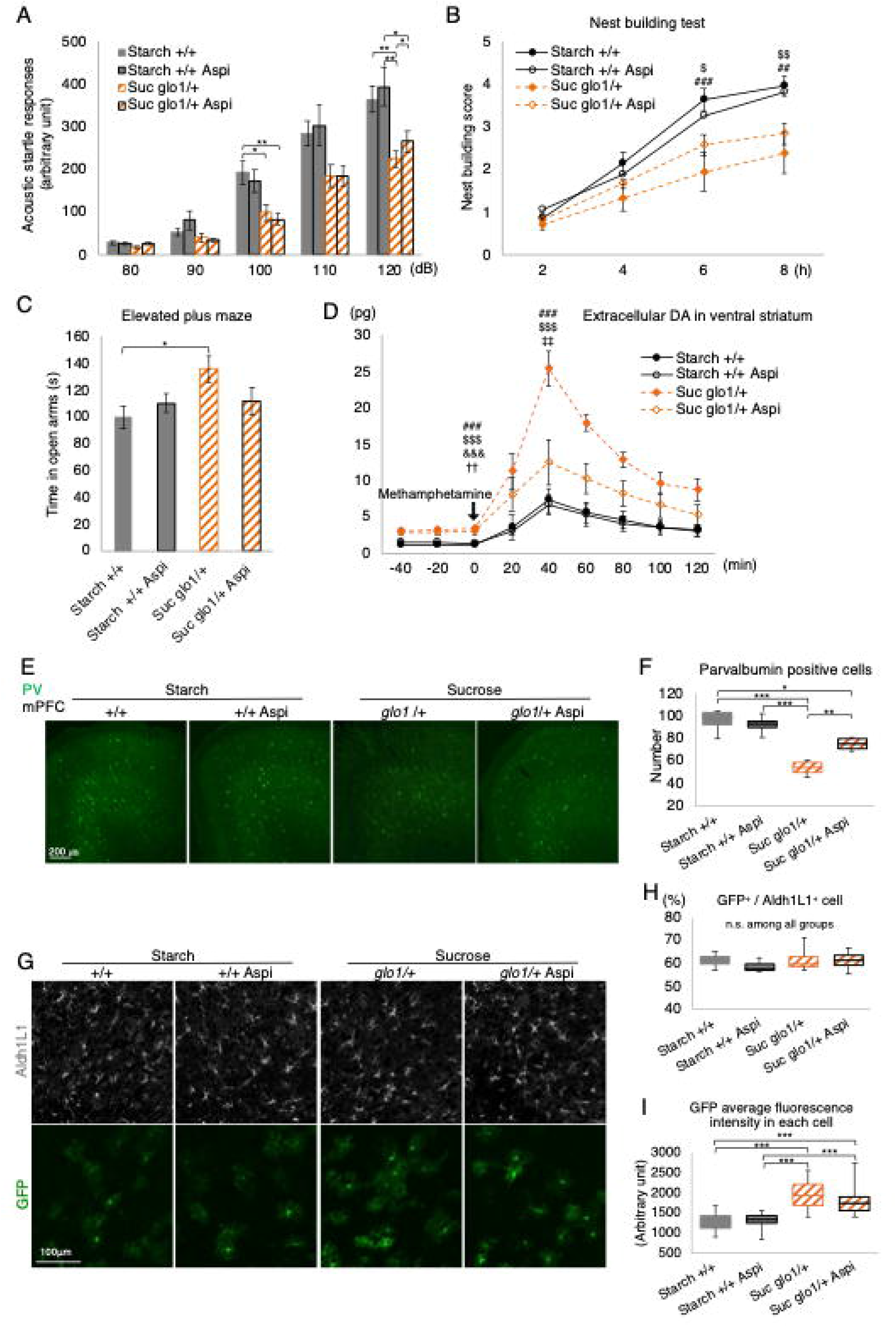
Protective effects of aspirin against the development of psychiatric phenotypes in G × E mice. (**A–C**) Effects of aspirin treatment on the acoustic startle response (**A**), nest-building skill (**B**), elevated plus maze test performance (**C**) and dopamine concentration (**D**). (**E**) Parvalbumin (PV) Immunohistochemistry in the medial prefrontal cortex (mPFC). (**F**) Number of PV-positive cells in the mPFC (n = 4 mice per group). (**G**) Immunohistological images of GFP-positive astrocytes and ALDH1L1in the mPFC region. (**H**) Percentage of GFP-positive cells per total ALDH1L1-positive cells in each image in (**G**). (**I**) Mean fluorescence GFP intensities of 10 randomly selected cells per image in (**G**) (from four independent mice). In **(B)**, post hoc Tukey‘s multiple comparisons test of groups at specific time points, ^###^*p* < 0.001, ^##^*p* < 0.01 for Starch +/+ vs. Suc *Glo1*/+ Aspi, ^$$^*p* < 0.01, ^$^*p* < for Starch +/+ Aspi vs. Suc *Glo1*/+ Aspi. In **(D)**, post hoc Bonferroni’s multiple comparisons test of groups at specific time points (0 min and 40 min), ^###^*p* < 0.001 for Suc *Glo1*/+ vs. Starch +/+, ^$$$^*p* < 0.001 for Suc *Glo1*/+ vs. Starch +/+ Aspi, ^&&&^*p* < 0.001 for Suc *Glo1*/+ Aspi vs. Starch +/+, ^††^*p* < 0.01 for Suc *Glo1*/+ Aspi vs. SucG*glo1*/+, ^‡‡^*p* < 0.01 for Suc *Glo1*/+ Aspi vs. Starch +/+ Aspi. All data are presented as mean ± SEM. \*\*\**p* < 0.001, \*\**p* < 0.01, \**p* < 0.05

**Supplementary Figure 8.**
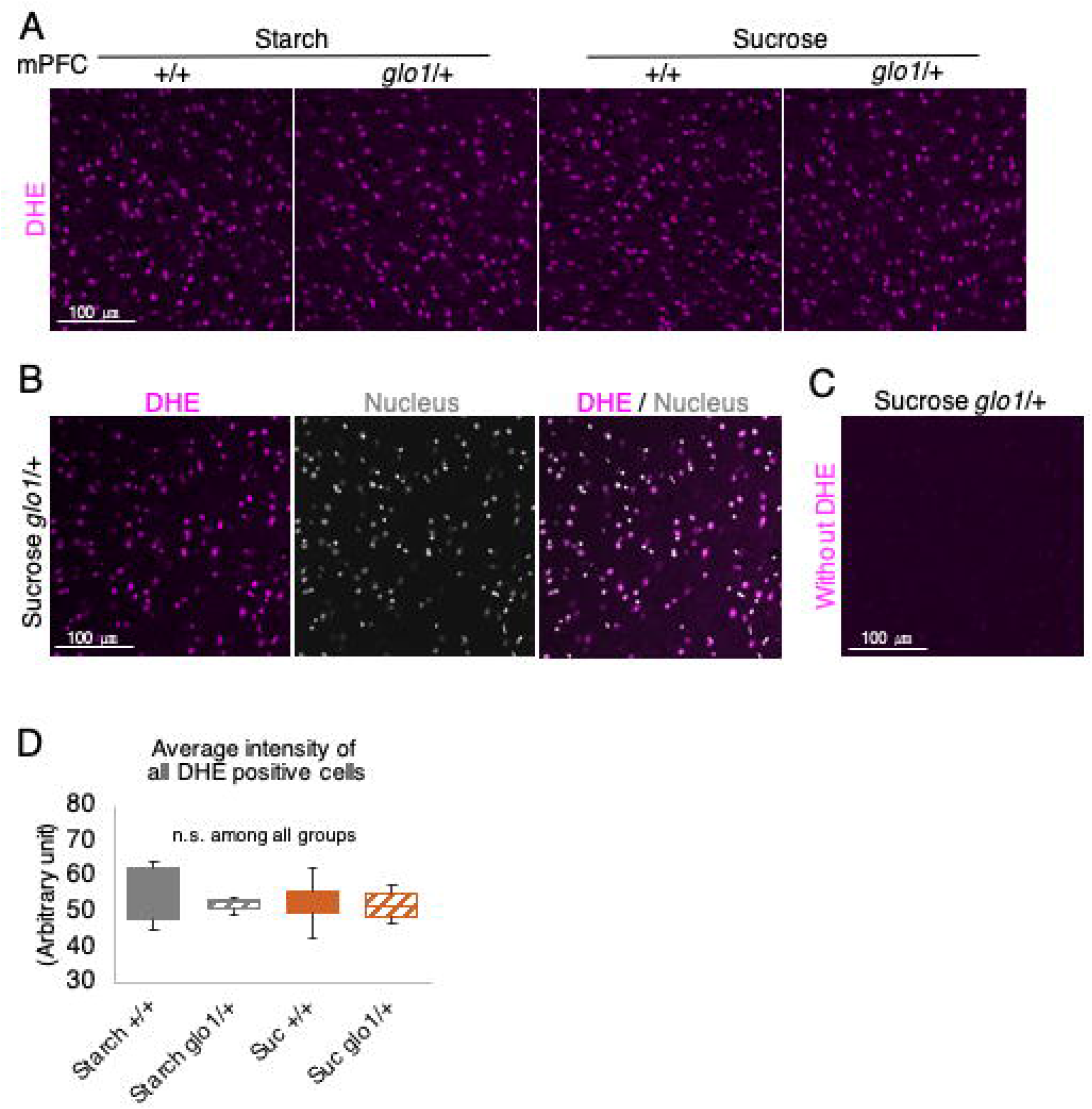
ROS detection in the four groups. (**A**) Reactive oxygen species (ROS) detection with *dihydroethidium* (*DHE*). (**B**) Merged DHE/Nucleus (TO-PRO 3) image. All cells are DHE positive, according to nucleus images. (**C**) ROS detection images without *DHE*. (**D**) Fluorescent intensity of DHE positive cells in (**A**). The mean DHE fluorescent intensity of each cell was measured in each section (n = 4 mice per group). All data are presented as mean ± SEM.

## Notes

### Competing Interest Statement

The authors have declared no competing interest.

